# Viral vector eluting lenses for single-step targeted expression of genetically-encoded activity sensors for in vivo microendoscopic calcium imaging

**DOI:** 10.1101/2023.11.09.566491

**Authors:** Carolyn K. Jons, David Cheng, Changxin Dong, Emily L. Meany, Jonathan J. Nassi, Eric A. Appel

**Affiliations:** Department of Materials Science & Engineering, Stanford University, Stanford CA 94305, USA; Inscopix - A Bruker Company, 1212 Terra Bella Ave. Suite 200, Mountain View CA 94043, USA; Department of Bioengineering, Stanford University, Stanford CA 94305, USA; Department of Pediatrics (Endocrinology), Stanford University, Stanford CA 94305, USA; ChEM-H Institute, Stanford University, Stanford CA 94305, USA; Woods Institute for the Environment, Stanford University, Stanford CA 94305, USA

**Keywords:** Optogenetics, Drug Delivery, Intravital Imaging, Neuroimaging, Calcium Imaging, GRIN Lenses

## Abstract

Optical methods for studying the brain offer powerful approaches for understanding how neural activity underlies complex behavior. These methods typically rely on genetically encoded sensors and actuators to monitor and control neural activity. For microendoscopic calcium imaging, injection of a virus followed by implantation of a lens probe is required to express a calcium sensor and enable optical access to the target brain region. This two-step process poses several challenges, chief among them being the risks associated with mistargeting and/or misalignment between virus expression zone, lens probe and target brain region. Here, we engineer an adeno-associated virus (AAV)-eluting polymer coating for gradient refractive index (GRIN) lenses enabling expression of a genetically encoded calcium indicator (GCaMP) directly within the brain region of interest upon implantation of the lens. This approach requires only one surgical step and guarantees alignment between GCaMP expression and lens in the brain. Additionally, the slow virus release from these coatings increases the working time for surgical implantation, expanding the brain regions and species amenable to this approach. These enhanced capabilities should accelerate neuroscience research utilizing optical methods and advance our understanding of the neural circuit mechanisms underlying brain function and behavior in health and disease.

## 1. Introduction

Advancements in neuroscience have been significantly accelerated by optical methods that allow researchers to study neural activity and its relationship with animal behavior[1],[2],[3]. Microendoscopic calcium imaging with miniature microscopes is a powerful approach for studying the relationship between neural activity and behavior in freely behaving animals[4],[5],[6],[7],[8]. The technique relies on the use of genetically encoded calcium indicators, most commonly GCaMP, to monitor large-scale neural activity in specific anatomically and genetically defined cell populations[9],[10]. Adeno-associated virus (AAV) injections are widely used for expressing GCaMP in target brain regions[11],[12]. Following an AAV injection, a second surgical step is required for implantation of an optical device, most commonly a gradient refractive index (GRIN) lens, for monitoring fluorescence deep in the brain[5],[13],[14]. This two-step procedure results in reduced experimental success rates due to an increased incidence of tissue damage and difficulties in targeting both the virus and the optical lens to the same site within the brain (**Fig. 1a, Figure S1a,b**). These challenges limit throughput and scale for translational and preclinical studies[15].

**Figure 1.**
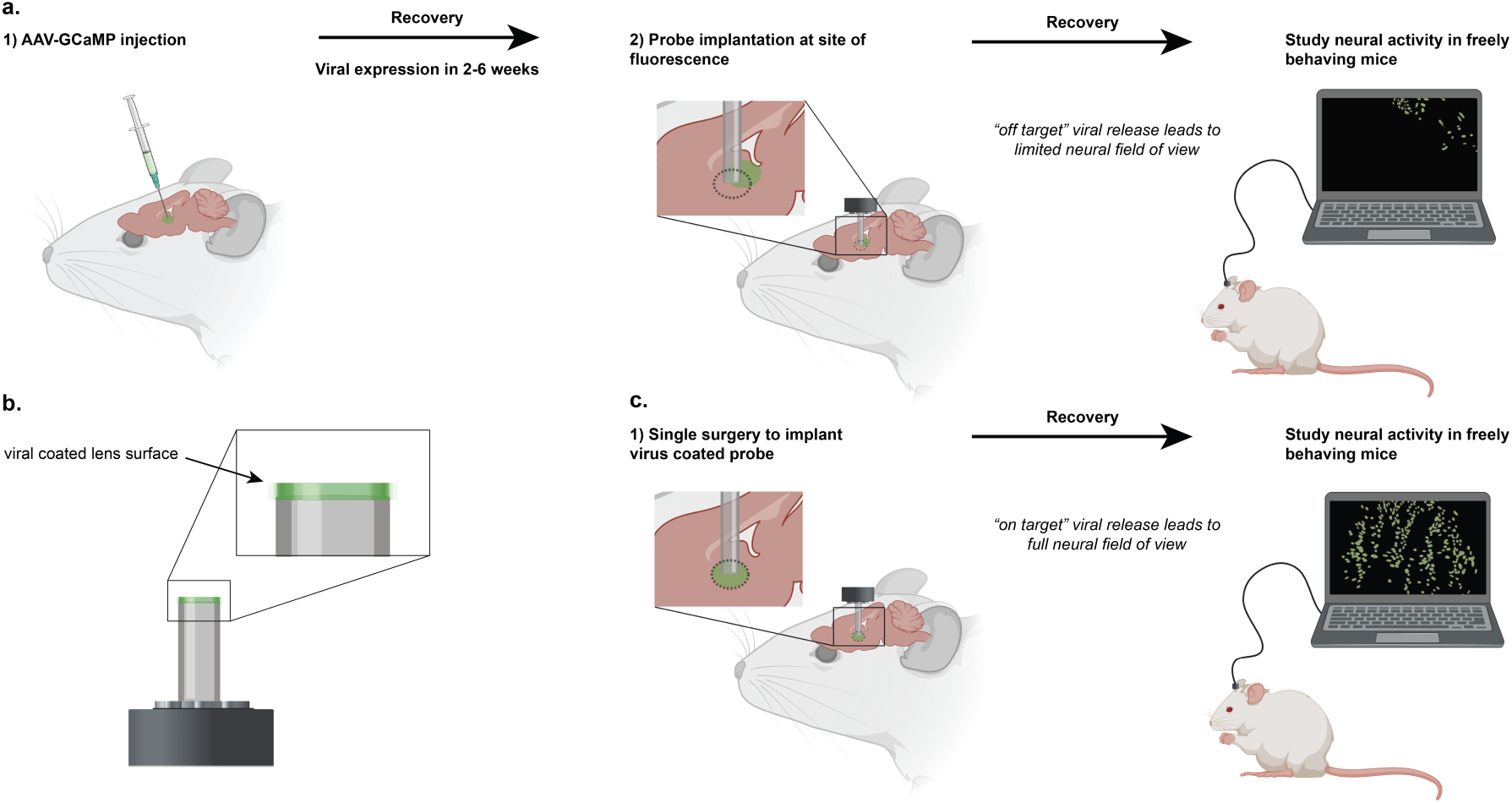
Delivery methods of AAV-GCaMP to study neural activity in freely behaving animals. **a)** Schematic illustrating traditional, two-step surgery process for separate virus injection and lens implantation, often resulting in misalignment and neural expression restricted to only part of the field of view. **b)** Lens coated with a polymeric material that encapsulates AAV-GCaMP and enables delayed and prolonged AAV release. **c)** Schematic illustrating how a lens coated with virus can enable neural expression over the full field of view following only a single surgical step. Created with BioRender.com

Prior research has sought to address these challenges by suspending AAV in silk fibroin films, which are then applied to the end of a GRIN lens for direct viral delivery to the imaging site of interest in a single surgical step[15]. These films are formed from drying aqueous silk fibroin derived from the cocoon of *Bombyx mori*[16],[17]. This method is inherently limited by the high solubility of silk fibroin, which results in immediate dissipation of the virus upon contact with cerebrospinal fluid. To circumvent this limitation, surgeons must rapidly implant the coated lens, resulting in the application of tensile and compressive stress to brain tissue, potentially causing tissue damage during the procedure. Moreover, the high solubility of silk films hinders its application in delivering viruses in more ventral brain regions and in species with larger brains than mice (e.g., rats and non-human primates). Apart from limitations in viral delivery efficiency, silk fibroin coatings are expensive, challenging to manufacture, and have a short shelf life[18]. To address these challenges and expand the number of supported imaging applications, it is imperative to develop a technological solution that stabilizes viruses and delays viral release during implantation, allowing for more working time and expanding the technology’s use in deeper brain regions and across different species. Delaying viral release upon implantation is critical to ensure precise targeting and effectiveness of the imaging approach.

Polymers have previously been shown as a promising material class for AAV delivery[19, 20]. Here, we engineered an AAV-eluting polymer coating for GRIN lenses that enables delayed and extended local delivery of AAV directly to the target brain tissue following implantation of the lens (**Fig. 1b**). These lens coatings reduce the number of required surgeries to one and guarantee alignment between GCaMP expression and lens placement in the brain (**Fig. 1c**). To achieve this controlled and localized viral delivery, we screened a library of polysaccharides with varied molecular weights and chemical compositions to identify coating materials that facilitate slow AAV diffusion and appropriately delayed and prolonged AAV exposure. Bench assays were developed and implemented to identify promising coating candidates based on cargo release kinetics, swelling behavior, biocompatibility, and ability to stabilize virus infectivity during storage. Lead candidates were then evaluated in-vivo through implantation of GRIN lenses coated with AAV-GCaMP-eluting polymer films into the dorsal medial striatum and medial prefrontal cortex (mPFC) of mice and rats. These studies demonstrated that lead polymer-coated lenses consistently provided high-cell-count fields-of-view with calcium transient kinetics and signal-to-noise ratios supporting high quality in-vivo calcium imaging. Critically, by engineering a polymer coating with delayed viral release, we increased lens implantation working time to enable microendoscopic imaging in additional brain regions (e.g., mPFC) and species (e.g., rats) that were not previously possible with other similar approaches (e.g., silk films). Overall, we report a versatile and practical solution to enable single-step delivery of genetically encoded sensors and actuators in specific brain regions to improve monitoring and control of neural activity.

## 2. Results and Discussion

Polysaccharides are a promising class of material for controlled therapeutic delivery because they are relatively inexpensive, available at scale, offer diverse chemistries and molecular weights, and demonstrate good biocompatibility and biodegradability[21],[22],[23]. A library of cellulose-based polysaccharides of varied molecular weights and functional groups were screened in-vitro to identify compositions providing cargo diffusion kinetics appropriate to slow the release of virus in the brain. Polymer solutions were dried in well plates to create defined films, and the release of incorporated virus-sized fluorescent dextran (MW ∼ 2,000 kDa) was quantified using a plate reader with fluorescent measurement capabilities (**Fig. 2a**). The higher molecular weight varieties of hydroxypropyl methyl cellulose (HPMC; MW ∼ 86 kDa) and sodium carboxymethyl cellulose (CMC; MW ∼ 700kDa) polymers were found to be promising candidates, demonstrating delayed cargo release within the first 20 minutes and nearly complete dextran release within 4 hours (**Fig. 2b**). Alternative cellulose candidates showed more rapid dextran release, and a sucrose control demonstrated full release of dextran within 5 minutes.

**Figure 2.**
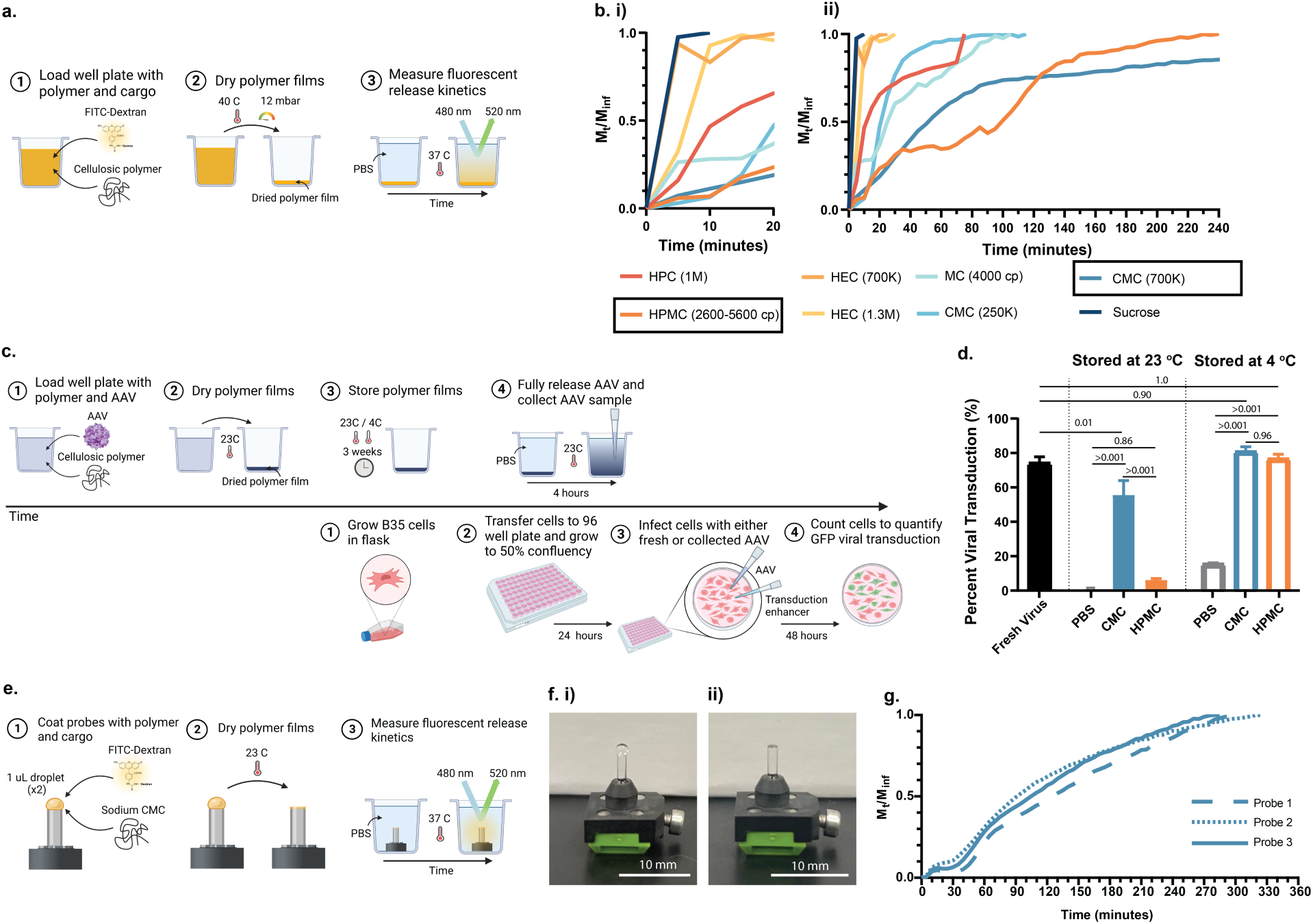
High-throughput screen to identify polymeric materials with promising viral release kinetics and viral viability. **a)** Method for preparing polymeric films in well plates to characterize dextran release kinetics. **b)** Rate of dextran release from thin films into PBS over i) 20 minutes and ii) 4 hours. **c)** Method for quantifying viral viability following encapsulation and storage in a polymeric film. **d)** Comparative viral transduction of fresh virus and virus released from polymeric films following three weeks of storage. **e)** Method for applying polymer films encapsulating dextran to GRIN lenses and measuring corresponding release kinetics. **f)** Illustration of 1 μL of polymer solution applied to a grin lens **i)** before and **ii)** after drying into a film. **g)** Rate of dextran release into PBS from CMC polymer films applied to GRIN lenses. Values reported are means ± SD of *n* = 3 per group. Statistical significance values are *p* values obtained from a Tukey HSD test. Created with BioRender.com

HPMC and CMC polymer candidates were further screened for their biocompatibility and ability to stabilize viral infectivity when dried into films. We conducted a viral transduction assay comparing the percent of B35 rat neuroblastoma cells transduced by AAV-GCaMP when AAV was stored and released from HPMC or CMC polymer films as compared to a freshly thawed AAV control stock (**Fig. 2c**). When films were stored in refrigerator conditions (4 °C) for 3 weeks, AAV released from HPMC and CMC films showed comparable viral transduction efficiency to that of the fresh AAV control. Films stored at room temperature (23 °C) for 3 weeks showed reduced viral transduction efficiency of AAV from both polymers (with respect to fresh AAV control), but transduction efficiency of AAV released from CMC films was statistically greater than that of AAV released from HPMC. Regardless of storage condition, AAV released from dried PBS showed a significant reduction in viral transduction efficiency, suggesting that encapsulation in a polymeric film prevents the loss of viral infectivity (**Fig. 2d**). As the viral transduction efficiency of AAV stored at room temperature was most improved when encapsulated in CMC (700kDa), this polysaccharide was selected as the lead polymer candidate for future development.

To confirm favorable release kinetics upon application to GRIN lenses, a 2 wt% solution of CMC containing virus-sized fluorescent dextran was applied to the surface of a GRIN lens using a pipette and allowed to dry (**Fig. 2e, f**). Using scanning electron microscopy (SEM), we determined that 4 μL of solution applied onto a 1 mm diameter lens resulted in a ∼160 μm thick dextran-encapsulating polymeric film. Cargo release kinetics from polymer films coated onto GRIN lenses were evaluated using a fluorescent plate reader assay. Again, delayed cargo release was observed over the first 20 minutes and nearly complete cargo release and film dissolution occurred within 6 hours (**Fig. 2g**).

CMC films were then assessed for their ability to enhance delivery of AAV to the brain in mice. GRIN lenses coated with 4 μL of 2 wt% CMC containing AAV encoding GCaMP were implanted into mouse dorsal medial striatum, and miniscope imaging sessions were conducted as animals freely explored an open field. Initial implant attempts resulted in dynamic flashes of diffuse fluorescence, but no clearly identifiable cells. Post-mortem histology in these cases revealed evidence of significant swelling of the polymeric coating in the brain (**Figure S2**). We hypothesized this swelling occurred shortly following initial contact with the brain and its constituent fluids (e.g., cerebrospinal fluid and blood), and that this swelling caused GCaMP-expressing neurons to lie beyond the focal plane of the imaging field of view.

To reduce the swelling of the polymer films when first contacting the brain, we reduced the film thickness from 160 μm to 80 μm by reducing the volume of polymer solution applied onto the GRIN lenses. To further reduce swelling, we also evaluated the blending of CMC with trehalose (Tre), a non-glycemic, high Tg sugar that is often used as a stabilizer in biopharmaceuticals[24],[25]. We hypothesized that by blending CMC with Tre while maintaining the same total solids content, we would reduce the entanglement of the polymers within the film, thereby reducing entropic swelling[26]. Polymeric films were formed from solutions comprising 2 wt% solids with different ratios of CMC to Tre. The notation “CMCX:TreY” indicates the volume fraction of 2 wt% stock solutions of CMC and Tre that were combined to form the final polymer solutions (i.e., the fraction of solids that is CMC and Tre (**Figure S3**). Film swelling and rate of cargo release was quantified with a film swelling assay whereby 80 μm thick polymer coatings were prepared on GRIN lenses, removed from the lenses, and then placed on top of an agarose brain mimic. Photographs were taken of the films over time with a fluorescent microscope to capture film swelling and cargo release (**Fig. 3a, b**). We observed decreased swelling and increased rate of cargo release at higher Tre contents (**Fig. 3c, d**). Increased rate of cargo release with higher Tre content likely results from faster film dissolution due to the higher water solubility of Tre as compared to CMC.

**Figure 3.**
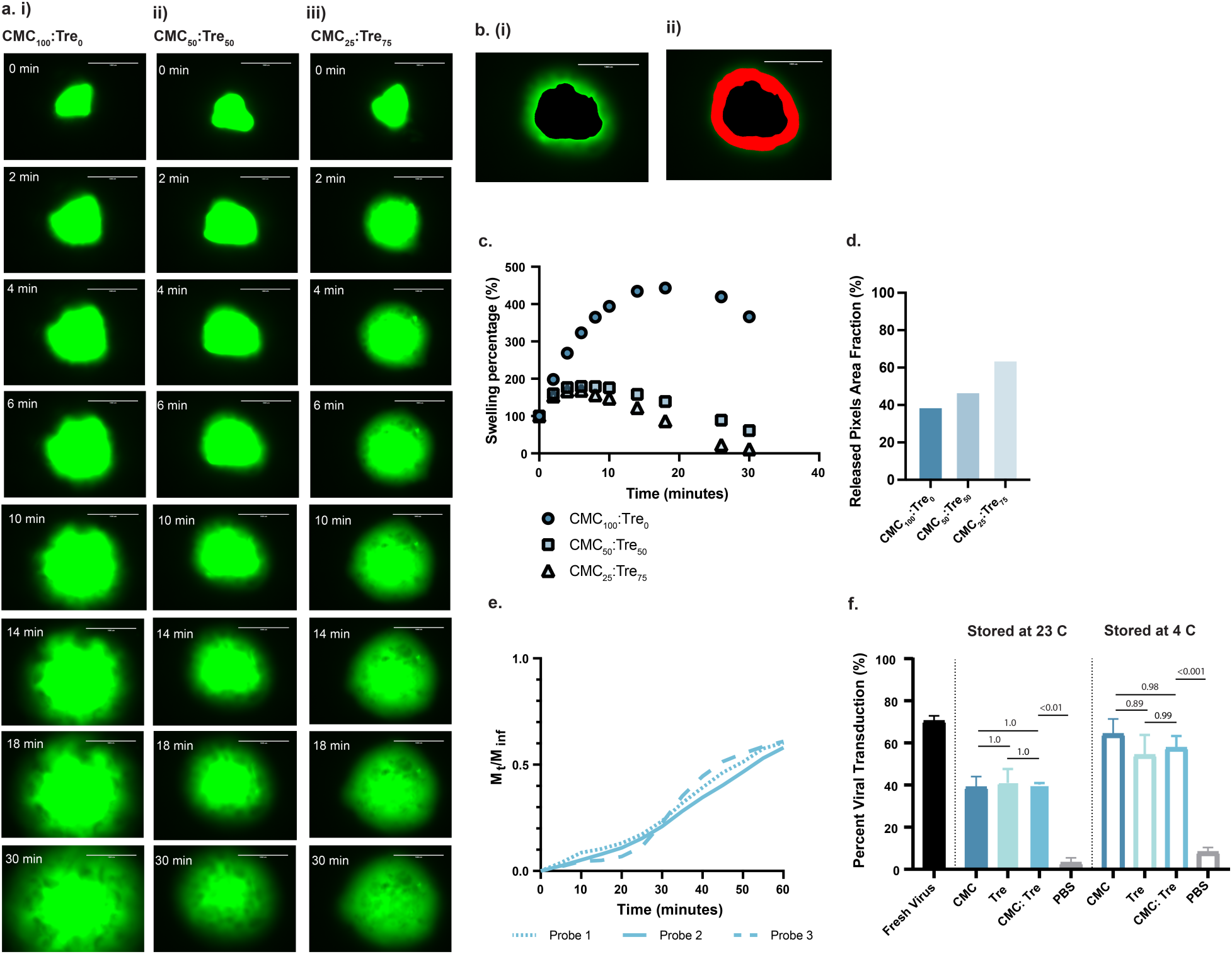
Trehalose content dictates CMC polymer film swelling and cargo release rates. **a)** Fluorescent microscopy images showing swelling and release kinetics of dextran loaded CMC: Tre films at a ratio of **(i)** CMC100:Tre0, **(ii)** CMC50:Tre50, and **(iii)** CMC25:Tre75. **b)** Representative ImageJ selection of film edges and released pixel area for quantification of swelling and relative release. Representative image corresponds to CMC50:Tre50 at 10 minutes. **c)** Quantification of swelling of CMC: Tre films. **d)** Quantification of released pixel area for CMC: Tre films at 10 minutes. **e)** Rate of dextran release into PBS from CMC30:Tre70 polymer films applied to GRIN lenses. **f)** Comparative viral transduction of fresh virus and virus released from CMC, trehalose, and CMC50:Tre50 films following three weeks of storage. Values reported in plot c and d are obtained from the film swelling images in 3a and are n=1. Values reported in plot f are means ± SD of *n* = 3 per group. Statistical significance values are *p* values obtained from a Tukey HSD test.

To mitigate the film swelling in the brain, thinner films (thickness ∼ 80 μm) with high Tre content (CMC30:Tre70) were selected as the leading film candidates in future experiments. Fluorescent microscopy illustrated uniform loading of dextran throughout the polymer film (**Figure S4**). Implantation into an agar brain mimic demonstrated CMC30:Tre70 films remained durable and adhered to the GRIN lens throughout the implantation process (**Figure S5**). Plate reader release assays with fluorescent dextran again confirmed promising release kinetics, whereby delayed dextran release was observed within the first 20 minutes (10 ± 3% dextran release) (**Fig. 3e**) and full dextran release and complete film dissolution occurred within 3 hours. Additionally, a viral transduction assay confirmed that films containing both CMC and Tre stabilize AAVs stored in polymer films. Films with both CMC and Tre show comparable viral transduction efficiency (p=0.98) to those only containing CMC following 3 weeks of storage in either refrigerator or room temperature conditions (**Fig. 3f**).

**Figure 4.**
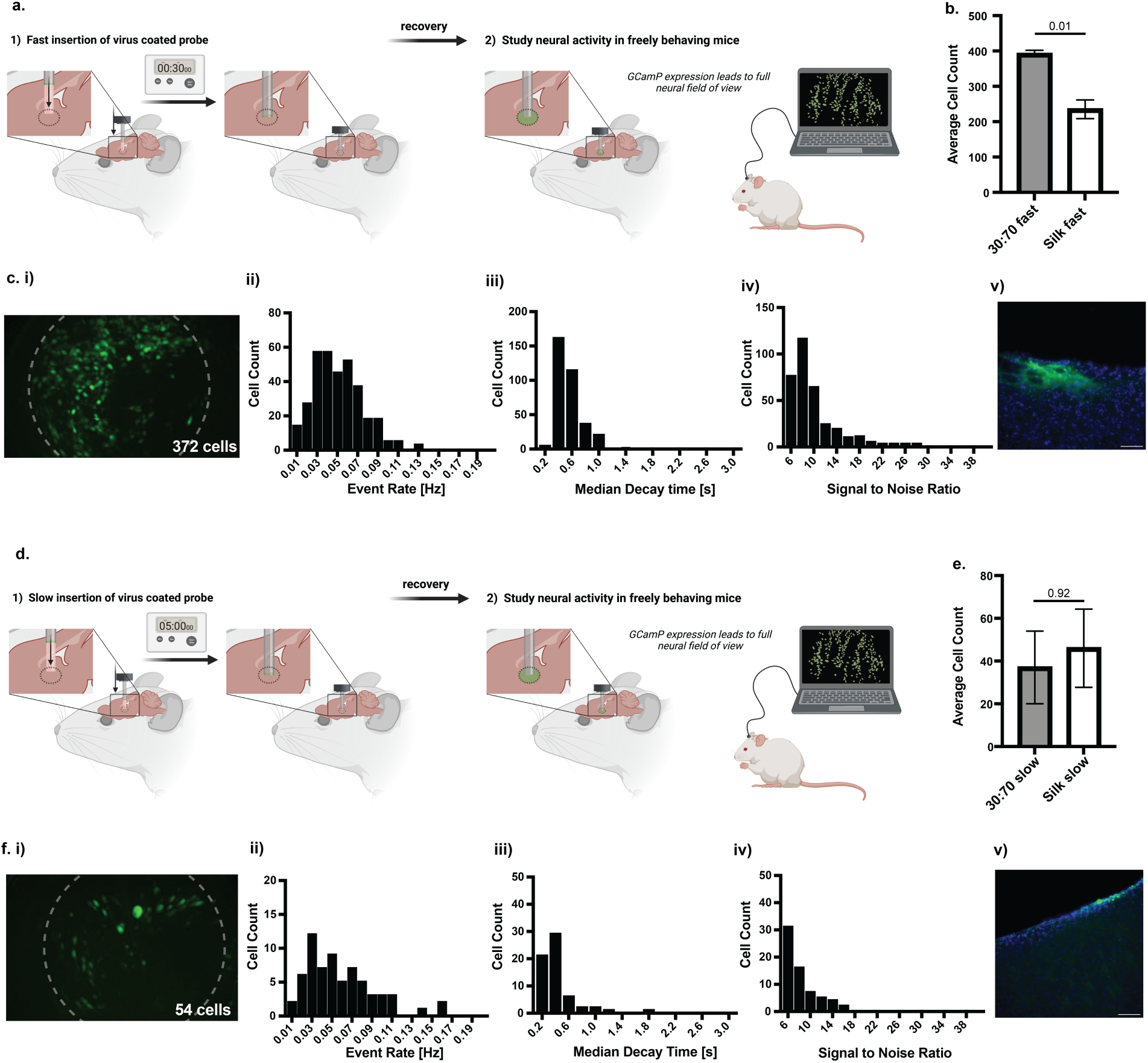
CMC30:Tre70 films result in viral expression in-vivo, but cell counts and field of view are dependent on implant procedure protocol. **a)** Schematic showing a fast implantation of a GRIN lens. **b)** Fast implant average cell count for CMC30:Tre70 films compared to silk fibroin control. **c)** Representative **i)** max projection image (GCaMP cells in green), histogram of neuronal calcium transient **ii)** median event rate, **iii)** median decay time, and **iv)** median signal to noise ratio and **v)** post-mortem histology for fast CMC30:Tre70 implants. **d)** Schematic showing a slow implantation of a GRIN lens. **e)** Slow implant average cell count for CMC30:Tre70 films compared to silk fibroin control. **f)** Representative **i)** max projection image, histogram of neuronal calcium transient **ii)** median event rate, **iii)** median decay time, and **iv)** median signal to noise ratio, and **v)** post-mortem histology for slow CMC30:Tre70 implants. Histology scale bars are 100 μm. Values reported in plots b and e are means ± SE of *n* = 2-3 per group. Statistical significance values are *p* values obtained from a two-tailed T test. Created with BioRender.com

**Figure 5.**
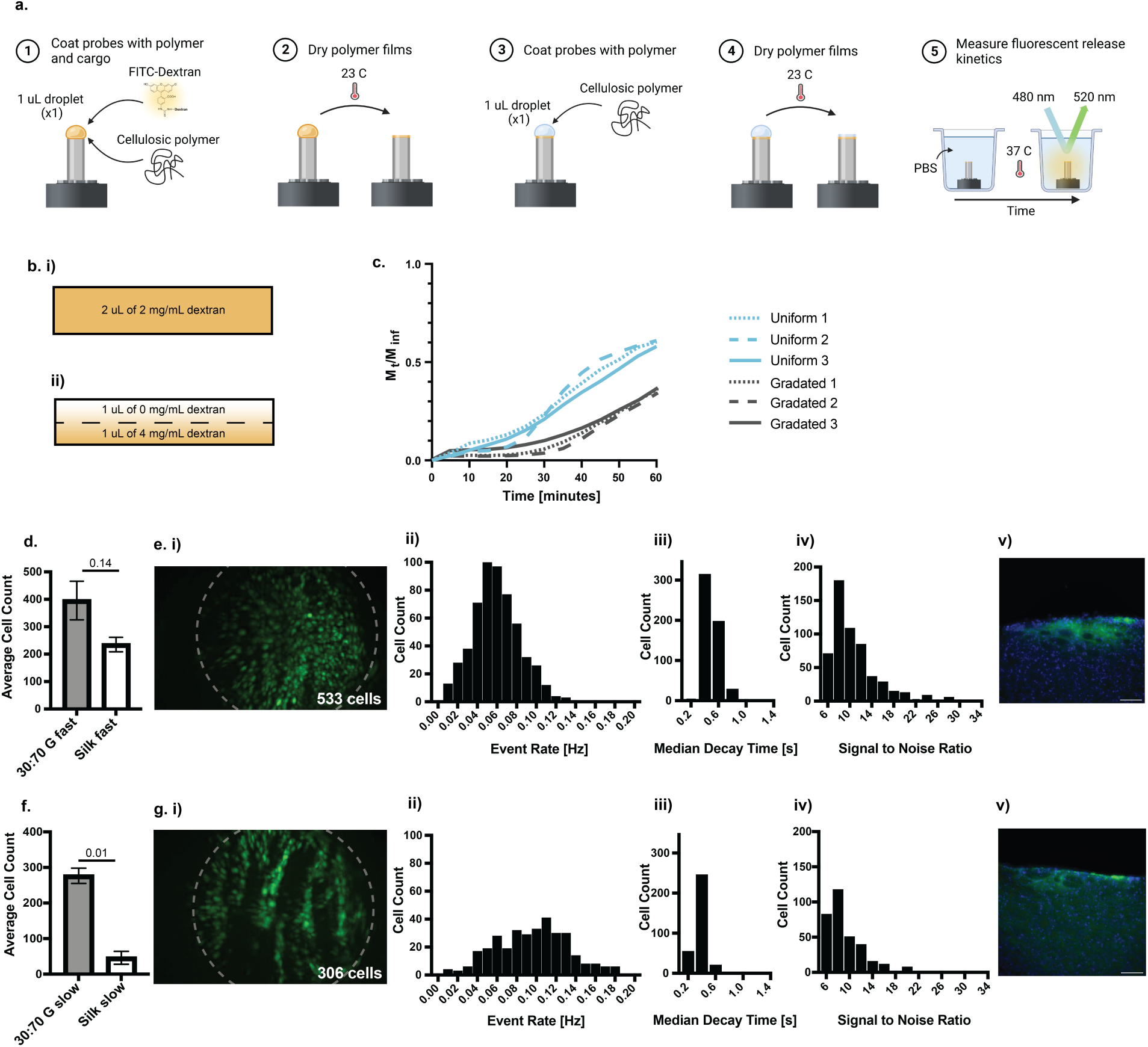
Gradated coatings allow for delayed viral release resulting in in-vivo viral expression independent of implant procedure protocol. **a)** Method for applying gradated polymer films encapsulating dextran to GRIN lenses and measuring corresponding release kinetics. **b)** Schematic of **i)** uniform dextran loading and **ii)** gradated dextran loading. **c)** Rate of dextran release into PBS from gradated and uniform CMC30:Tre70 polymer films applied to GRIN lenses. **d)** Fast implant average cell count for CMC30:Tre70G films compared to silk fibroin control. **e)** Representative **i)** max projection image (GCaMP cells in green), histogram of neuronal calcium transient **ii)** median event rate, **iii)** median decay time, and **iv)** median signal to noise ratio, and **v)** post-mortem histology for fast CMC30:Tre70-G implants. **f)** Slow implant average cell count for CMC30:Tre70-G films compared to silk fibroin control. **g)** Representative **i)** max projection image (GCaMP cells in green), histogram of neuronal calcium transient **ii)** median event rate, **iii)** median decay time, and **iv)** median signal to noise ratio, and **v)** post-mortem histology for slow CMC30:Tre70-G implants. Histology scale bars are 100 μm. Values reported in plots d and f are means ± SEM of *n* = 3-5 per group. Statistical significance values are *p* values obtained from a two-tailed T test. Created with BioRender.com

The new polymer film formulation was then screened for in-vivo functionality. GRIN lenses coated with 2 μL of 2 wt% CMC30:Tre70 solutions containing AAV-GCaMP were implanted into the mouse dorsal medial striatum, and miniscope imaging sessions were again conducted as animals freely explored an open field. To evaluate the impact of implantation speed on efficacy of GCaMP expression and neuronal imaging, we performed two versions of the implant procedure that varied in speed of device placement. In the fast implant scenario, a tract was first made in the brain, and the optical probe was inserted to a final depth of 2.4 mm within 30 seconds (**Fig. 4a**). This fast implant procedure provides minimal time for virus release, ensuring the majority of encapsulated virus reaches the brain region of interest. This implant procedure has previously been shown to be successful when a commercial silk fibroin coating is used to deliver AAV-GCaMP (**Figure S6**). In the slow implant procedure, brain probes were instead inserted slowly to a final depth of 2.4 mm over the course of 5 minutes (**Fig. 4d**). This slow insertion process was designed to mimic more challenging implant conditions associated with deeper insertion depths and higher prevalence of fluids of other brain regions and larger species. We anticipated the extended film contact with blood and cerebrospinal fluids in the brain arising from the slow insertion would increase the likelihood of undesirable, premature virus release before the probe reaches the brain region of interest, thereby challenging the capabilities of our new polymer coatings to sufficiently delay virus release. Current commercial silk fibroin coatings do not result in adequate neuronal GCaMP expression in this slow implant scenario due to rapid virus release (**Figure S7**).

**Figure 6.**
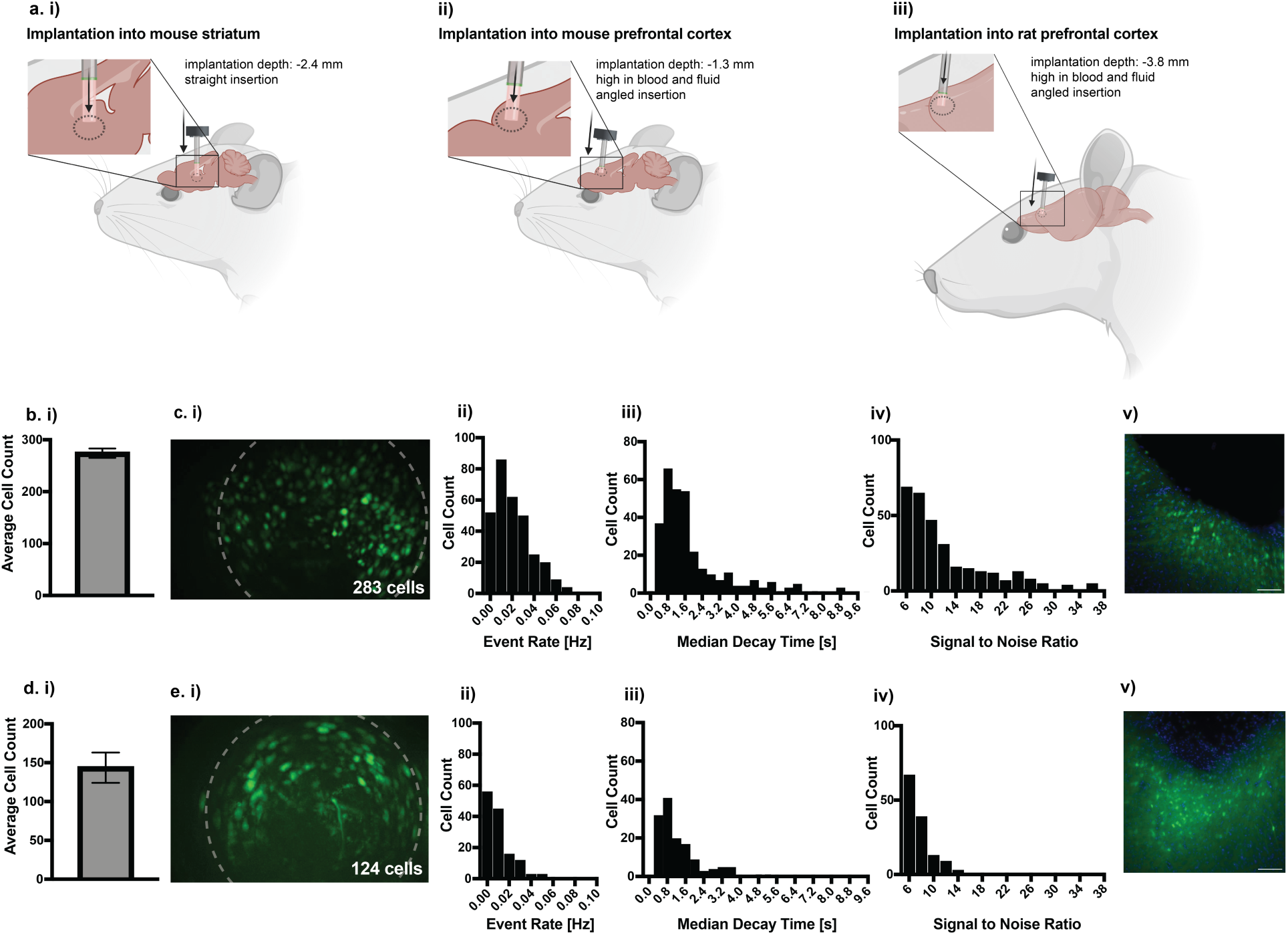
Gradated coatings allow for delayed viral release resulting in in-vivo viral expression in new brain regions and in new species. Schematic illustrating **a. i)** implant into mouse striatum, **ii)** implant into mouse mPFC, and **iii)** implant into rat mPFC. **b)** Average cell count for CMC30:Tre70-G films implanted into the mouse mPFC. **c)** Representative **i)** max projection image (GCaMP cells in green), histogram of neuronal calcium transient **ii)** median event rate, **iii)** median decay time, and **iv)** median signal to noise ratio, and **v)** post-mortem histology for CMC30:Tre70-G implants in the mouse mPFC. **d)** Average cell count for CMC30:Tre70-G films implanted into the rat mPFC. **e)** Representative **i)** max projection image (GCaMP cells in green), histogram of neuronal calcium transient **ii)** median event rate, **iii)** median decay time, and **iv)** median signal to noise ratio, and **v)** post-mortem histology for slow CMC30:Tre70-G implants in the rat mPFC. Histology scale bars are 100 μm. Values reported in plots b and d are means ± SEM of *n* = 2 per group. Created with BioRender.com

Fast implant studies were conducted with probes coated with 80 μm thick coatings of (i) CMC30:Tre70, (ii) CMC50:Tre50, or (iii) silk fibroin controls comprising AAV encoding GCaMP. Probes with CMC:Tre based coatings dramatically outperformed those with commercial silk fibroin controls, resulting in high counts of labeled cells within the field of view, favorable neuronal event rates, favorable decay times, and low signal-to-noise ratios (**Fig. 4b, Figure S8, Figure S9, Table S1**). In these studies, imaging field of views with CMC50:Tre50 coated devices showed greater translational movement than CMC30:Tre70 coated devices, likely due to an increased degree of swelling associated with the higher CMC content in the films. Moreover, post-mortem histological analysis showed a viral expression zone for CMC30:Tre70 coated devices within the working distance of the lens (**Fig. 4c**).

While fast implants resulted in promising neuronal expression, slow implants with these 80 μm thick CMC30:Tre70 coatings resulted in low cell counts (**Fig. 4e**), even though neuronal event rates, decay times, and signal-to-noise ratios remained promising (**Fig. 4f**). We hypothesized that the AAV-GCaMP release from the films is not sufficiently delayed in-vivo to survive the slow probe insertion process. To further delay the rate of release of virus and improve the performance of our coating in a slow implant protocol, we sought to generate bilayer gradated CMC30:Tre70 coatings (CMC30:Tre70-G). Gradated coatings were applied to GRIN lenses by first applying a 1 uL droplet of 2 wt% CMC30:Tre70 solution loaded with cargo at twice the concentrations used previously, allowing this film to dry completely, and then applying a second 1 μL droplet of 2 wt% CMC30:Tre70 solution containing no virus so that it formed a sacrificial polymer layer to delay virus release (**Fig 5a**). The kinetics of cargo release from the gradated film was compared to that of a uniform film formed from application of a 2 μL droplet of polymer solution (**Fig 5b**). Release kinetics from polymer coated GRIN lenses were confirmed using the fluorescent plate reader assays described above. Gradated films exhibited completely negligible cargo release over the first 20 minutes (4 ± 2% dextran release) (**Fig 5c**) and slow release after that, but still showed complete cargo release and film dissolution within 5 hours.

GRIN lenses coated with this gradated coating were implanted into mouse dorsal medial striatum, and miniscope imaging sessions were again conducted as animals freely explored an open field. Gradated coatings allowed for delayed release of virus and targeted viral delivery in both fast and slow implant conditions. With both fast and slow implantations, 80 μm thick CMC30:Tre70-G coatings dramatically outperformed silk fibroin control coatings. For both implant procedures, CMC30:Tre70-G coatings resulted in an increased cell count within the field of view, improved neuronal calcium transient event rates, improved decay times, and lower signal-to-noise ratios compared to silk fibroin controls (**Fig. 5d,f; Figure S10, Table S1, Video S1, Video S2**). Moreover, post-mortem histology showed a clear viral expression zone within the working distance of the lens (**Fig. 5e, g, Figure S11**). These observations suggest the CMC30:Tre70-G coatings are more robust to the conditions of implantation in the brain and produce more consistent, higher yield fields of view for in-vivo imaging studies.

For commercial applications of coated GRIN lens devices, coatings must exhibit suitable shelf-life for distribution and storage over several months. We found the CMC30:Tre70-G coated GRIN lenses maintain AAV transfection efficiency following 16 weeks of storage at 4 °C (**Figure S12**). This in-vivo data corroborates prior viral transduction data suggesting CMC and trehalose aid in viral stabilization in a dried state (**Fig. 3f**).

Following development of a polymeric coating delaying viral release and enabling viral delivery in more challenging implant scenarios, GRIN lenses coated with CMC30:Tre70-G films containing AAV encoding GCaMP were implanted into the mPFC of mice and rats. While the mPFC of a mouse requires a shallower implantation depth (1.3 mm), implantation in this region typically must overcome more blood and fluid and requires an angled insertion to avoid midline vasculature. Implantation in the mPFC of a rat typically must overcome even more blood and fluid, and again requires an angled insertion. Furthermore, the depth of implantation (3.8 mm) is more ventral than both dorsal striatum and mPFC in mice. (**Fig. 6a**). For these reasons, implantation in the mPFC of mice and rats was previously not possible with silk fibroin film (**Figure S13**).

CMC30:Tre70-G coatings in the mPFC of both mice and rats resulted in high cell counts within the field of view (**Fig. 6b,d; Figure S13, Video S3, Video S4**). Implants additionally showed favorable neuronal event rates, decay times, and signal-to-noise ratios, and post-mortem histology showed a viral expression zone within the working distance of the lens (**Fig. 6c,e**).

## 3. Outlook

This work details the development of a simply, versatile, and practical polysaccharide-based coating that can be applied to the imaging face of a GRIN lens for locoregional delivery of genetically encoded calcium indicators, such as GCaMP, to specific brain regions upon implantation of the lenses in the tissue of interest. *In-vivo* validation of our approach demonstrates that the polymer coated lenses consistently provide high cell count fields of view with calcium transient kinetics and signal-to-noise ratio supporting high quality in-vivo calcium imaging. Importantly, the polymer coating not only stabilizes the AAVs prior to delivery, allowing for shelf-stable manufacturing, but also slows down their release *in-vivo*, extending the time available for lens implantation and enabling the exploration of deeper brain regions in various species. These capabilities represent an important technical advance that promises to accelerate breakthrough discoveries and enable higher throughput and scale for translational and preclinical neuroscience research. While this study demonstrates the coating of GRIN lenses and implantation into the brains of mice and rats, we anticipate this technology to be widely applicable to alternative imaging modalities and additional brain regions and species. The combination of optical tools with our novel polymer coating represents a crucial step forward in neuroscience research, facilitating a deeper exploration of neural dynamics and paving the way for transformative discoveries in the field of biomaterials and neural engineering.

## 4. Methods

### 4.1 Materials

Hydroxypropyl Cellulose (MW∼1,000,000), Methyl Cellulose (viscosity 4,000 cP), Sodium Carboxymethyl Cellulose (MW∼700,000), Sodium Carboxymethyl Cellulose (MW∼250,000), Hydroxypropylmethyl Cellulose (viscosity 2,600-5,600 cP; MW∼86,000), Hydroxyethyl Cellulose (MW∼1,300,000), Hydroxyethyl Cellulose (MW∼700,000), Trehalose dihydrate (>99.0%), Sucrose, FITC-dextran (MW∼2,000,000), silk fibroin, and Poly(L-lysine) were obtained from Sigma-Adrich and used as received. Sterile Phosphate buffered saline (Gibco, pH 7.4) was obtained from Thermo Fisher Scientific and used as received. Cell culturing supplies including DMEM, FBS, and Pen/Strep were obtained from Cytivia. Etoposide and DMSO were obtained from Sigma-Aldrich. AAV DJ of pAV-CMV-GFP virus and AAV1-CaMKII-GCaMP6m virus were obtained from Vigene Biosciences and stored at -80 °C prior to use.

### 4.2 In-vitro characterization

#### High throughput polymer screen

To estimate the kinetics of viral release from the polymeric films, films that contain an AAV-sized fluorescent dextran are prepared[27],[28], and release rate of this dextran into PBS is measured using a plate reader. FITC-dextran (MW 2,000,000) is used as a model cargo. A dextran stock solution is prepared at a concentration of 20 mg mL^-1^ in PBS. Then 25 μL of this stock solution is combined with 1 mL of 2 wt% polymer solution in PBS to achieve a final concentration of 0.5 mg mL^-1^ dextran. 200 μL of polymer-dextran solution is added to each well, and polymer-dextran solutions are dried using a Genovac (EZ2-Elite, Low BP method). After films are dried, wells are filled with 200 μL of PBS using a multi channel pipette, and the 96 well plate is covered with an optically transparent plate seal. The rate of dextran release from the polymer films is measured using a plate reader (Synergy H1 Microplate reader, 23 °C, excitation: 480 nm, emission 520 nm). Complete release was seen after approximately 4 hours. Kinetic release curves are normalized to the highest fluorescence value.

#### Viral transduction assay

Viral transduction experiments were performed to quantify viral viability in polymeric films following storage. Rat neuroblastoma B35 cells were obtained as a frozen culture from the Stanford University Neuroscience Gene Vector and Virus Core. Cells were cultured in a T75 flask at 37 °C with 5% CO2 in DMEM supplemented with high glucose, L-glutamine, sodium pyruvate (Cytiva, SH30243.FS), 10% heat inactivated fetal bovine serum (Cytiva, SH30396.03HI), and 1% penicillin (100 U mL^-1^)/streptomycin (100 μg mL^-1^) (Cytiva, SV30010). Cells were split 2-3 times prior to experimental use after growing to 70-80% confluency.

Viral transduction experiments require plating cells in a cell-adherent 96 well plate. Cells were first grown to 80% confluency in a T75 flask in media containing DMEM, 10% FBS, and 1% pen-strep. Media was removed from the T75 flask using an aspirator, and adherent cells were rinsed with 2 to 3 mL of warmed 1X PBS. PBS was then aspirated. To lift cells from the T75 flask, 2 to 3 mL of warmed trypsin was added and left for 2 to 3 minutes. To stop the trypsin action, 6 mL of warmed media was added. The media-trypsin mixture was transferred to a 10 mL centrifuge tube, and cells were pelleted by spinning at 125 G for 7 minutes. Following spinning, the supernatant was removed, and the cell pellet was redispersed in 1 mL of warmed media. Cell count of this stock cell solution was quantified using a LUNA-FL Dual Fluorescence Cell counter, fluorescent cell counting slides, and an acridine orange-propidium iodide dye following procedures recommended by Logos Biosystems. The stock cell solution was diluted with warmed, fresh media to a concentration of 1 x 10^5^ live-cells mL^-1^. 200 μLs of the 1 x 10^5^ cell mL^-1^ solution were added to each well of the cell-adherent 96 well plate (20,000 cells/well), and cells were incubated for 24 hours or until 50% confluent as observed through bright field microscopy.

After cells reach 50% confluence, AAV is introduced to each well. To improve viral transduction, media was carefully removed with a P200 pipette and replaced with warmed media containing the transduction enhancer etoposide. Etoposide was initially dissolved at 25mM in DMSO, and diluted 1000X with fresh media (ex. 20 mL media, 20 μL etoposide stock). AAV DJ of pAV-CMV-GFP of the desired concentration was then added to each well. For most experiments, this was 0.1 μL of 7.28 x 10^13^ GC mL^-1^ AAV (7.28 x 10^9^ GC/well). Cells with added virus were incubated for 24 to 48 hours.

Following 24 to 48-hour incubation, viral transduction efficiency was quantified. Media was carefully aspirated from each well. Wells were then rinsed with 40 μL of warmed PBS, and PBS was aspirated. To lift cells from the wells, 30 μL of warmed trypsin was added and left for 2 to 3 minutes. To stop the trypsin action, 100 μL of warmed media was added. Media and cells were mixed gently with a pipette, and transduction efficiency was quantified using a LUNA-FL Dual Fluorescence Cell counter and low-fluorescent cell counting slides following the GFP expression procedures recommended by Logos Biosystems. Transduction efficiency was quantified as the number of GFP transfected cells divided by total cell count.

#### Quantification of viral viability in films

To measure viral viability in the polymeric film, fresh virus was added to polymer solutions, dried into a film, and released into PBS. Viral transduction efficiency of virus released from films was compared to that of fresh virus. Films were formed by combining 1 μL of fresh virus with 200 μL of 2 wt% solutions of HPMC, CMC, Trehalose, a 50:50 blend of CMC and Trehalose dissolved in PBS, or PBS alone. These polymer solutions were dried into thin films in a 96 well plate. After films were dried, they were stored either at 23 °C or 4 °C for 3 weeks. Following storage, 200 μL of PBS was added to each well, and films were allowed to fully dissolve. Following complete dissolution after 4 hours, a 20 μL sample (containing 0.1 μL of virus) was taken from each well. The viral transduction efficiencies of these viral samples were compared to that of fresh virus in the viral transduction assay.

#### Lens coating process (uniform and gradated)

To estimate the kinetics of viral release from the polymeric films, probes are coated with polymer containing a comparable-size fluorescent dextran, and release rate of this dextran into PBS is measured using a plate reader. Fluorescence isothiocyanate dextran (MW 2,000,000) is used as a model cargo.

#### Uniform, ∼160 μm thick film

A dextran stock solution is prepared at a concentration of 20 mg mL^-1^ in PBS. Then 10 μL of this stock solution is combined with 200 μL of 2 wt% polymer solution in PBS to achieve a final concentration of 1 mg mL^-1^ dextran. Probes are coated with four 1 μL droplets of the polymer-dextran solution (0.004 mg dextran/ probe), and the polymer film is given time to dry between each droplet application.

#### Uniform, ∼80 μm thick film

A dextran stock solution is prepared at a concentration of 20 mg mL^-1^ in PBS. Then 20 μL of this stock solution is combined with 200 μL of 2 wt% polymer solution in PBS to achieve a final concentration of 2 mg mL^-1^ dextran. Probes are coated with two 1 μL droplets of the polymer-dextran solution (0.004 mg dextran/ probe), and the polymer film is given time to dry between each droplet application.

#### Gradated, ∼80 μm thick film

A dextran stock solution is prepared at a concentration of 20 mg mL^-1^ in PBS. Then 20 μL of this stock solution is combined with 100 μL of 2 wt % polymer solution in PBS to achieve a final concentration of 4 mg mL^-1^ dextran. Probes are first coated with a 1 μL droplet of the polymer-dextran solution (0.004 mg dextran/ probe) and films are allowed to dry. A 1 μL droplet of 2 wt% polymer solution without dextran is then added on top of the dried film, and the film is again given time to dry.

#### Kinetics release assay from coated probes

Probes are adhered to the base of a 96 well plate using a small amount of vacuum grease. Vacuum grease is not in contact with the polymer film. Wells are filled with 250 μL of PBS using a multi-channel pipette, and the 96 well plate is covered with an optically transparent plate seal. The rate of dextran release from the polymer films is measured using a plate reader (Synergy H1 Microplate reader, 37 °C, orbital shaking, excitation: 480 nm, emission 520 nm). Complete release was seen for all samples after 4 to 6 hours. Kinetic release curves are normalized to the highest fluorescence value.

#### Quantification of film swelling

A dextran stock solution is prepared at a concentration of 20 mg mL^-1^ in PBS. Then 20 μL of this stock solution is combined with 200 μL of 2 wt % CMC-trehalose solution in PBS to achieve a final concentration of 2 mg mL^-1^ dextran. Probes are coated with two 1 μL droplets of the polymer-dextran solution (0.004 mg dextran/ probe), and the polymer film is given time to dry between each droplet application. Films are manually removed from the lens surface. Film swelling is quantified when the film is in contact with an agarose brain mimic. The agarose brain mimic is formed by adding agarose to distilled water at 0.6 wt%. The solution is heated while stirring to 85 °C until agarose is fully dissolved, and the agar is then cooled to form a gel in clear petri dishes. Films are placed on the surface of the agar, and film swelling and dextran release over 30 minutes is imaged with a fluorescent microscope. Images are analyzed in ImageJ to quantify the change in area as films swell as well as fraction of dextran release over time. In this analysis, we first compare the area of the solid green in the original film to the area of solid green in films at subsequent timepoints to quantify film swelling. We also compare the area of diffuse green pixels (as compared to total area which includes solid green regions plus diffuse pixels) to quantify fraction of released pixels.

#### Quantification of Uniformity of Cargo Loading

GRIN lens coatings were prepared with a dextran concentration of 1 mg mL^-1^. Polymer films were carefully removed from the GRIN lens and imaged using a fluorescent microscope. Uniformity of dextran loading was quantified in ImageJ by taking the mean grey value (average fluorescent intensity) of a rectangle 1/10^th^ the diameter of the polymer film.

#### Stability of Film During Implantation

GRIN lens coatings were prepared with a dextran concentration of 1 mg mL^-1^. An agrose brain mimic was created as described above. Polymer films on GRIN lenses were first imaged using an optical and fluorescent microscope. GRIN lenses were then inserted and removed from the agrose brain mimic and imaged again with an optical and fluorescent microscope.

#### SEM Imaging

Films were characterized by scanning electron microscopy (SEM). Samples were grounded to a 90-degree aluminum pin stub using double-sided conductive copper tape. A 5.0 nm thick layer of pure gold was deposited onto the samples using a Leica ACE600 Vacuum system. SEM analysis was performed using the FEI Magellan 400 XHR Scanning Electron Microscope at 5.00 kV and high vacuum in field-free mode.

#### Fourier Transform Infrared Spectroscopy

Fourier transform infrared spectroscopy (FT-IR) was performed on each sample using an attenuated transmission reflectance (ATR) setup on a Nicolet-iS50 spectrometer at the Stanford Soft and Hybrid Materials Facility (SMF). The lyophilized powder was loaded on the diamond window of the ATR setup, and reflectance was observed using a KBr beamsplitter for wavenumbers 400 – 5000 cm^-1^. Sample data were collected in duplicate.

### 4.3 In-vivo methodology

#### Animal subjects

All procedures were conducted in accordance with the Guide for the Care and Use of Laboratory Animals, as adopted by the National Institutes for Health, and with approval of the NASA Institutional Animal Care and Use Committee. Adult (25 - 30 g) C57BL/6J mice and 180-280g Sprague-Dawley rats were separately group housed until surgery with ad libitum access to food and water. Mice were maintained on a reverse 12 hour light cycle (lights off at 8:00 a.m.).

#### Viral Constructs

Purified and concentrated adeno-associated virus serotype 1 encoding the calcium indicator GCaMP6m driven by the CaMK2 promoter (AAV1-CaMKII-GCaMP6m) were packaged by Vigene Biosciences at titers 1.59 x 10^13^ genome copies (GCs) mL^-1^.

#### Viral Injection Surgeries

For all surgeries, mice (n=2) were anesthetized with 1.5–2.0% isofluorane mixed with 1 L per min of oxygen and placed in a stereotaxic frame (Kopf Instruments, Tujunga, California). Body temperature was maintained at 37 °C with a heating pad (40-90-2-07, FHC, Bowdoin, Maine). Mice and rats received subcutaneous injections of ketoprofen (2.5 mg kg^-1^) and carprofen (2.5 mg kg^-1^).

A vertical incision was made to expose the skull, and any residual tissue was cleared so that bregma and lambda were clearly visible. A small burr hole was made over the site of injection (Mouse Striatum: +1.0AP, ±1.5ML, -2.4DV, Mouse PFC: +2.0AP, ±0.3ML, -1.6DV). Using a hamilton syringe equipped with a 33G needle (35G for Mouse PFC), 1 μL of virus was drawn up into the syringe. The syringe was then attached onto the stereotax arm, held in place by a UMP3 Ultramicropump (WPI, Sarasota, FL) and the needle tip zeroed over the surface of the brain. The needle was then lowered through the burr hole to its respective injection depth, and virus injected at a rate of 100 nL min^-1^ (80 nL min^-1^ for Mouse PFC) for 500 nL total volume. A post infusion time of 5 minutes was allotted before removing the needle out of the tissue, at a rate of 50-100 μm sec^-1^. The incision over the skull was sutured and the animal was returned to its home cage, on a heating pad, to recover and wake before being placed back into the housing room, with 72 hour ad libitum access to 80 mg/ 200 mL acetaminophen in water. GRIN lens implantation occurred after 2 weeks and followed protocols described below.

#### Preparation of CMC:Trehalose coatings for in-vivo implantation

Polymer solutions were dissolved at 2 wt% in sterile PBS. After dissolution, polymer solutions were sterile filtered using sterile syringes and 0.22 μm syringe filters. The dissolved polymer solutions were combined with a viral stock of AAV1.Camk2a.GCaMP6m.WPRE.SV40 (1.59 x 10^13^ GC mL^-1^). These components were combined in the appropriate ratios to deliver a viral load of 5.28 x 10^9^ GC per probe. Prior to applying the polymer coating containing AAV, the distal, imaging lens face was cleaned with Sticklers Fiber Optic Splice and Connector Cleaner and Foamtec International Miraswab paddle tip swabs. The polymer solution was then applied to the surface of the GRIN lens using a P2 micropipette in 1 μL increments. Coating occurred at 4C, and films were allowed to fully dry before another polymer droplet was added to the surface. Completed films were stored in air-tight plastic boxes at 4 °C until implantation.

#### Preparation of silk fibroin coatings for in-vivo implantation

1x4mm GRIN lenses (Inscopix, PN:1050-004637) were coated at the distal, imaging lens face with Poly(L-lysine) (0.01%), silk fibroin (2.5%) and AAV1.Camk2a.GCaMP6m.WPRE.SV40 (5.28 x 10^9^ GCs). Probes were then placed into a humidity-controlled chamber overnight and allowed to air-dry the next morning, before being placed into membrane bound boxes and stored at 4 °C until time of implantation.

#### Coated Lens Storage Conditions

All coated lenses used were shipped and stored in membrane bound boxes at 4 °C until time of implantation. Long-term stability tests studied viral expression following 16 weeks of storage.

#### Lens Implantation Surgeries

For all surgeries, mice (n=35) and rats (n=2) were anesthetized with 1.5–2.0% isofluorane mixed with 1 L per min of oxygen and placed in a stereotaxic frame (Kopf Instruments, Tujunga, California). Body temperature was maintained at 37 °C with a heating pad (40-90-2-07, FHC, Bowdoin, Maine). Mice and rats received subcutaneous injections of ketoprofen (2.5 mg kg^-1^) and carprofen (2.5 mg kg^-1^).

A single skull screw (19010-10, Fine Science Tools, Foster City, California) was placed on the contralateral hemisphere, away from the lens coordinates. A craniotomy over the targeted brain region (Mouse Str; coordinates from bregma: +1.0 anterior, +1.65 lateral; Mouse PFC: +2.0AP, ±0.65ML) was performed and dura was carefully removed. The probe (either a ProView™ Express Probe (Silk fibroin formulation; Inscopix, Mountain View, California) or a ProView™ Integrated Lens (Inscopix PN: 1050-004367) coated with this new formulation) was removed from 4 °C and placed onto a threaded dummy microscope (Inscopix PN: 1050-003762) attached to a stereotax arm (Kopf Instruments, Tujunga, California) just prior to aspiration. Approximately 2.1 mm for Mouse Str and 1.2mm for Mouse PFC of tissue were aspirated using a 30-gauge blunt needle attached to a vacuum pump to allow for an entry path for the GRIN lens (approx. 1.0mm diameter x 4.0mm length). Saline was continuously applied during the aspiration to avoid drying of the tissue. Bleeding was controlled by sterile saline-soaked gelfoam. Immediately following aspiration, the lens was implanted to a depth of -2.4DV (Mouse PFC: -1.3DV at 10° angle) from the surface of the brain. Lenses were implanted at a rate of approximately 100um/sec for fast implantations, or less than 10um/sec for slow implantations. Rat PFC implantations occurred in a similar fashion. For the rat PFC (Coordinates: +3.0AP, ±0.85ML, -3.8DV at 5° angle), approximately 3.5 mm of tissue was aspirated before implantation of a 1.00 mm diameter x 9.0 mm length ProView™ Integrated Lens (Inscopix, PN:1050-004416) coated with the new formulation. Kwik-sil (World Precision Instruments, Sarasota, Florida) was applied around the craniotomy and base of lens to cover up any exposed brain tissue, and the remaining portions extending above the surface of the skull was fixed with metabond (Parkell, Edgewood, New York) and anchored by the skull screw. Imaging sessions ranged from 5 – 20 minutes in duration at 6 weeks post implantation at 20 frames per second (Inscopix Data Acquisition Software, Inscopix, Inc.). LED power (range: 0.2 – 1 mW) and gain (range: 2-8) sensor settings were adjusted for each recording session to approximately normalize the baseline range of fluorescent counts across all recording sessions within and across mice. After 8 weeks of study, all mice and rats in the study were euthanized and perfused transcardially with PBS followed by 4% paraformaldehyde (PFA), and the extracted brains were post-fixed in 4% PFA for approximately 12 hours.

#### In-Vivo Imaging Sessions

All imaging sessions were conducted within a (24in x 24in) open field chamber. Briefly, animals were scruffed, a miniature microscope docked onto their baseplate and placed into a chamber for freely behaving imaging. Microscope settings were fine tuned for each imaging session to obtain the appropriate field of view.

#### In-Vivo Imaging Analysis

All recording sessions were spatially down-sampled (2x), spatially cropped, corrected for rigid translational motion and normalized according to baseline fluorescence (Δf/f) and, finally, individual cells and their corresponding calcium traces were extracted using PCA/ICA (Inscopix Data Processing Software, Inscopix, Inc.). Cells with a minimum signal-to-noise ratio (SNR) greater than or equal to 5 and a spatial footprint consisting of a single connected component were included for further analysis. We quantified the median calcium event rise time, decay time, SNR and rate for each cell in each recording session and every subject. Events were detected as waveforms that crossed an adaptive threshold (5 times the median absolute deviation). Rise time for an event was calculated as half-height to peak event amplitude time. Event decay time was calculated as the time from event peak to half height. SNR was calculated as the median event peak amplitude divided by the median absolute deviation. Rates were calculated as numbers of calcium events per second over the entire recording.

#### Post-mortem histology

Brains were cut coronally on a vibratome with 50 μm thick sections. Sections were washed in PBS prior to preparation on slides for histological assessment and then DAPI stained (DAPI Fluoromount-G PN: 0100-20, Southern Biotech). Images were obtained on a Leica DMi8 inverted epi-fluorescent microscope.

### 4.4 Statistical methods

For in-vitro experiments, comparison between multiple groups was conducted with a Tukey HSD test, and values presented were means and standard deviations. Results were accepted as significant if *p* < 0.05. For in-vivo experiments, comparison between multiple groups was conducted with a Tukey HSD test and comparison between two groups was conducted with a two-tailed t-test in JMP. Values presented were means and standard errors. Results were accepted as significant if *p* < 0.05.

## 5. Acknowledgements

This research was financially supported by a gift from Inscopix, a Bruker Company, to the Stanford Neurophotonics Center. Part of this work was performed at the Stanford Nano Shared Facilities (SNSF), supported by the National Science Foundation under award ECCS-2026822. Part of the data was collected at the Stanford Soft and Hybrid Materials Facility (SMF) (supported by the NSF grant National Science Foundation grant ECCS1542152). C.K.J. is thankful for a National Science Foundation Graduate Research Fellowships. E.L.M. is grateful to the NIH Biotechnology Training Program (T32 GM008412) for funding. We are additionally grateful for the support of Javier Alcudia, and the Stanford Neuroscience Gene Vector and Virus Core for providing our B35 rat neuroblastoma cell line and recommendations for viral transduction assays. We thank Noah Eckman for his support with figure preparation.

## 6. Author Contributions

C.K.J., D.C., J.N. and E.A.A. conceived of the idea. C.K.J. and D.C. performed experiments and analyzed the data. C.D. and E.L.M. performed experiments. We thank Swathy Sampath Kumar, Mark Trulson, Keon Visscher and Oliver Miller for helpful discussions.

## 7. Declaration of Interests

D.C. and J.N. are full-time employees at Inscopix – A Bruker Company. All other authors declare that they have no competing interests.

## 8. Materials and Data Availability

All data needed to evaluate the conclusions in the paper are present in the paper and/or the Supplementary Materials. Further information and requests for resources or raw data should be directed to and will be fulfilled by the lead contact, Eric Appel.

## Supplementary Materials

Figures S1 to S13

Table S1

Videos S1 to S4

## Notes

### Summary of Updates

Updated PDF with images with improved resolution

