## Supplemental Material for "Viral vector eluting lenses for single-step targeted expression of genetically-encoded activity sensors for in vivo microendoscopic calcium imaging"

### Table of Contents

**Figure S1.** Mismatch between lens implantation location and viral expression zone…………..….3

**Figure S2.** CMC films result in swelling in-vivo……………………………………………..……….3

**Figure S3.** Optical, SEM images and FTIR of $CMC_{30}$:$Tre_{70}$ film and $CMC_{100}$ film……..............….4

**Figure S4.** Uniformity of cargo loading in $CMC_{30}$:$Tre_{70}$ film……………………………………......….5

**Figure S5.** Film adherence to GRIN lens during implantation process……………..……….…….6

**Figure S6.** Replicates of fast implants of commercial silk films…………………...……………..……7

**Figure S7.** Replicates of slow implants of commercial silk films………………………..………...…..8

**Figure S8.** Replicates of fast and slow implants of $CMC_{30}$:$Tre_{70}$ films………………………………9

**Figure S9.** Fast implants of $CMC_{50}$:$Tre_{50}$ films……………………………………….…..…..…….10

**Figure S10.** Replicates of fast and slow implants of $CMC_{30}$:$Tre_{70}$-G films…………..……………..11

**Figure S11.** Insertion traces of fast and slow implants………………………………….……..…….12

**Figure S12.** Fast implants of $CMC_{30}$:$Tre_{70}$-G films following 16 weeks of storage………..…….…13

**Figure S13.** Replicates of $CMC_{30}$:$Tre_{70}$-G films implanted into the mPFC…………...…..….……..14

**Table S1.** Summary of striatum and mPFC implants……………………………………………...15

**Video S1.** $CMC_{30}$:$Tre_{70}$-G fast implant into the mouse dorsal medial striatum……………....….17

**Video S2.** $CMC_{30}$:$Tre_{70}$-G slow implant into the mouse dorsal medial striatum...……...….….….17

**Video S3.** $CMC_{30}$:$Tre_{70}$-G implant into the mouse prefrontal cortex…………………......….…….17

**Video S4.** $CMC_{30}$:$Tre_{70}$-G implant into the rat prefrontal cortex………………...……...……….17

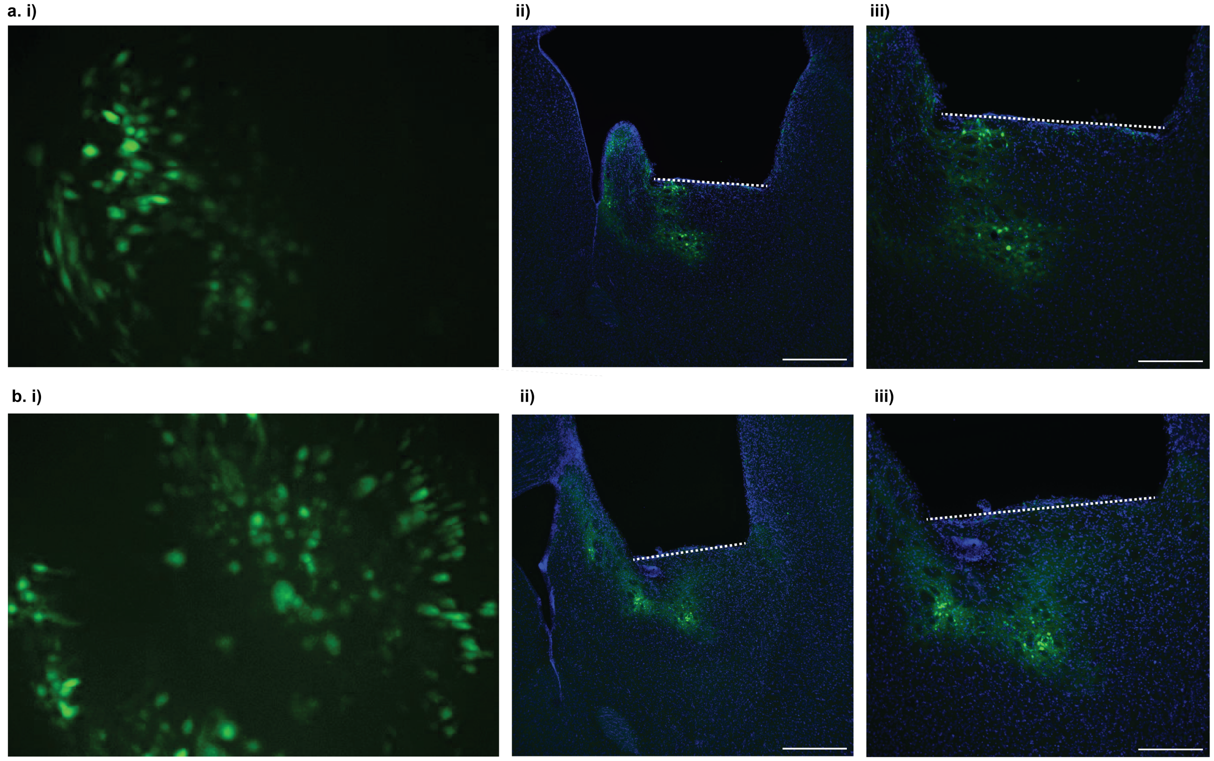


**Figure S1.** A two-step surgical method often results in mismatch between lens implantation location and viral expression zone. **a)** GCaMP expression zone is laterally offset in relationship to the implanted lens which results in a partial neuronal field of view during in-vivo imaging. **b)** GCaMP expression zone is ventrally offset in relationship to the implanted lens which results in poorer neuronal resolution during in-vivo imaging. Green is GCaMP expression and blue is DAPI-stained neuronal nuclei. Dashed white lines indicates position of imaging face of lens in the brain. Histology scale bars are **a. i)** 500 µm, **a. ii)** 250 µm, **b. i)** 500 µm, and **b. ii)** 250 µm.

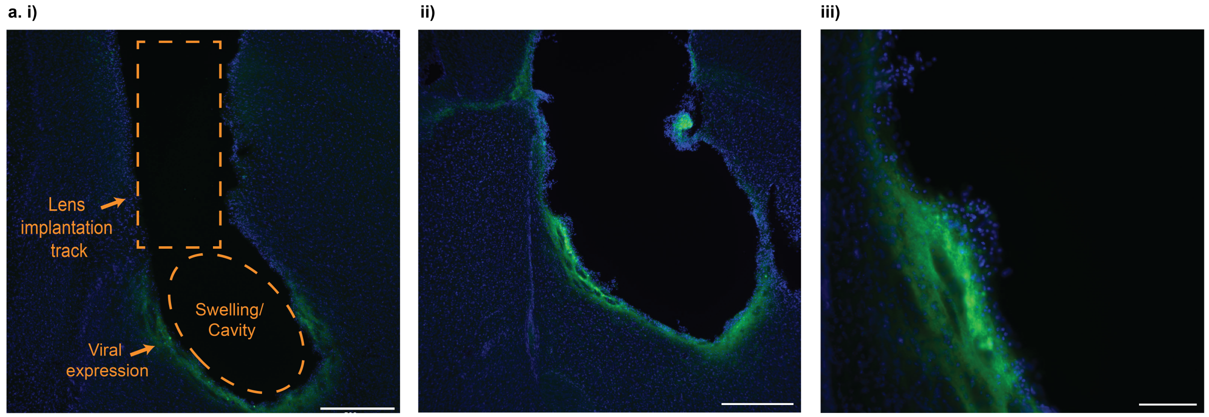


**Figure S2.** CMC polymer films result in viral expression, but significant film swelling results in tissue displacement and viral expression beyond the focal range of the lens. Histology with scale bars of **a)** 1 mm, **b)** 500 µm, **c)** 100 µm. GCaMP, green; DAPI-stained neuronal nuclei, blue.

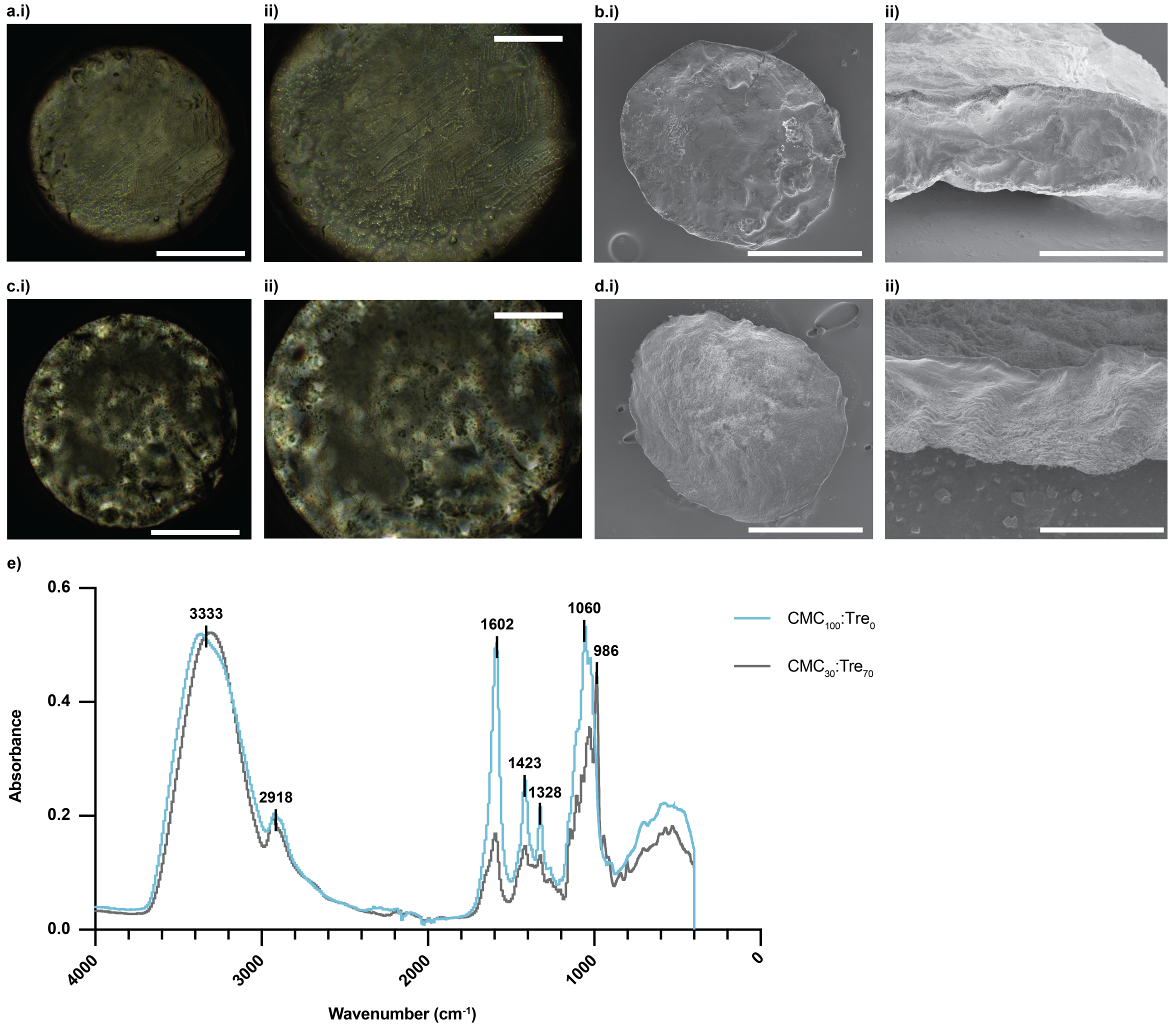


**Figure S3.** Optical and SEM images of $CMC_{30}$:$Tre_{70}$ film and $CMC_{100}$ film. **a)** Optical image of $CMC_{30}$:$Tre_{70}$ film on the GRIN lens surface with **i)** 400 μm scale bar and **ii)** 200 μm scale bar. **b. i)** Top surface of $CMC_{30}$:$Tre_{70}$ film removed from the GRIN lens surface with 400 μm scale bar and **ii)** cross-section of ~ 85 μm thick $CMC_{30}$:$Tre_{70}$ film with 100 μm scale bar. **c)** Optical image of $CMC_{100}$:$Tre_0$ film on the GRIN lens surface with **i)** 400 μm scale bar and **ii)** 200 μm scale bar. **d. i)** Top surface of $CMC_{100}$:$Tre_0$ film removed from the GRIN lens surface with 400 μm scale bar and **ii)** cross-section of ~ 85 μm thick $CMC_{100}$ with 100 μm scale bar. **e)** FTIR spectra for $CMC_{30}$:$Tre_{70}$ and $CMC_{100}$:$Tre_{0}$. Bands at 3333 and 2918 $cm^{-1}$ are due to -OH and C-H stretching vibrations, respectively. Absorption bands at 1602 and 1423 $cm^{-1}$ are assigned to the asymmetric and symmetric stretching of COO− group, respectively. Absorption bands at 1328 $cm^{-1}$result from –OH bending vibration. Absorption bands at 1060 $cm^{-1}$ are due to CH–O–CH2 stretching. Absorption bands at 986 $cm^{-1}$ are due to C-C stretching[1].

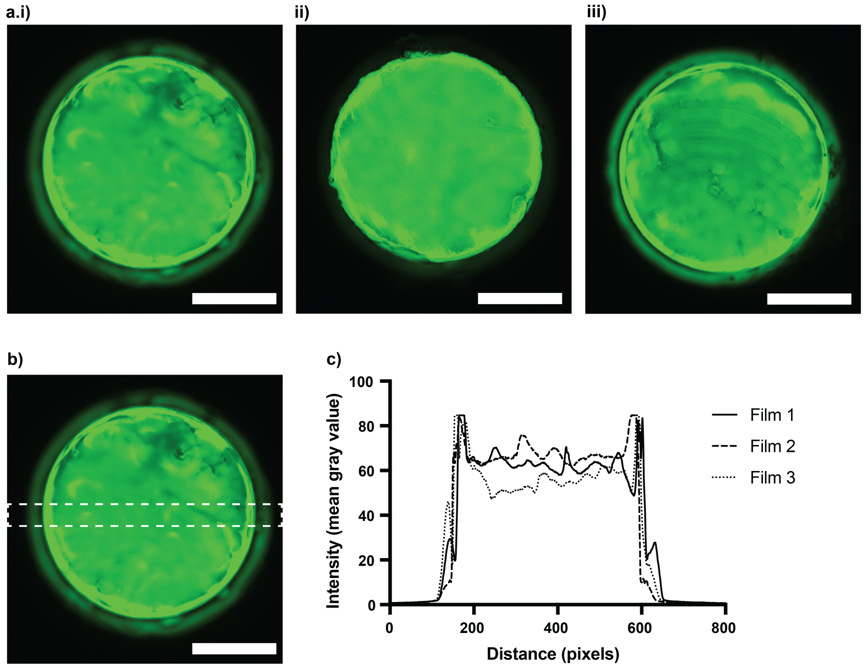


**Figure S4.** Uniformity of cargo loading in $CMC_{30}$:$Tre_{70}$ film. **a)** Fluorescent microscopy images of replicate **(i, ii, iii)** dextran-loaded $CMC_{30}$:$Tre_{70}$ films. **b)** Representative analysis region of average pixel intensity. **c)** Average pixel intensity across film surface.

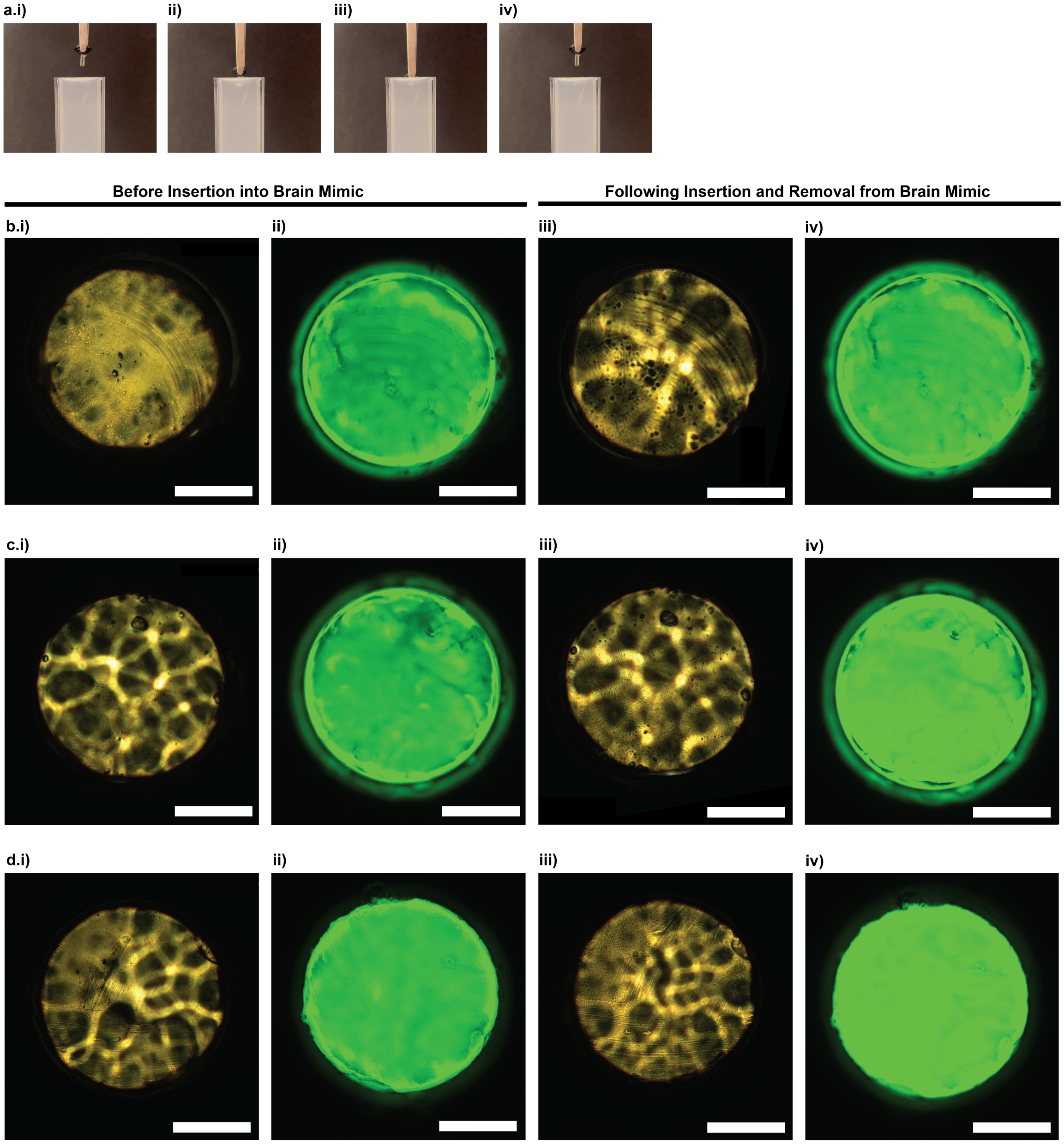


**Figure S5.** $CMC_{30}$:$Tre_{70}$ film adherence to GRIN lens during implantation process **a)** Pictures depicting implanting GRIN lens with $CMC_{30}$:$Tre_{70}$ film into agarose brain mimic showing **i)** pre-insertion, **ii)** partial insertion, **iii)** full insertion, and **iv)** removal. **b)** First replicate of optical **(i)** and fluorescent **(ii)** microscopy images of dextran-loaded $CMC_{30}$:$Tre_{70}$ film adhered to a GRIN lens before insertion into agarose brain mimic. Optical **(iii)** and fluorescent **(iv)** microscopy images of dextran-loaded $CMC_{30}$:$Tre_{70}$ film adhered to a GRIN lens following insertion and removal from agarose brain mimic. **c, d)** Second and third replicate of optical and fluorescent microscopy images of dextran-loaded $CMC_{30}$:$Tre_{70}$ film adhered to a GRIN lens before and after insertion and removal from an agarose brain mimic. Scale bars are 400 μm.

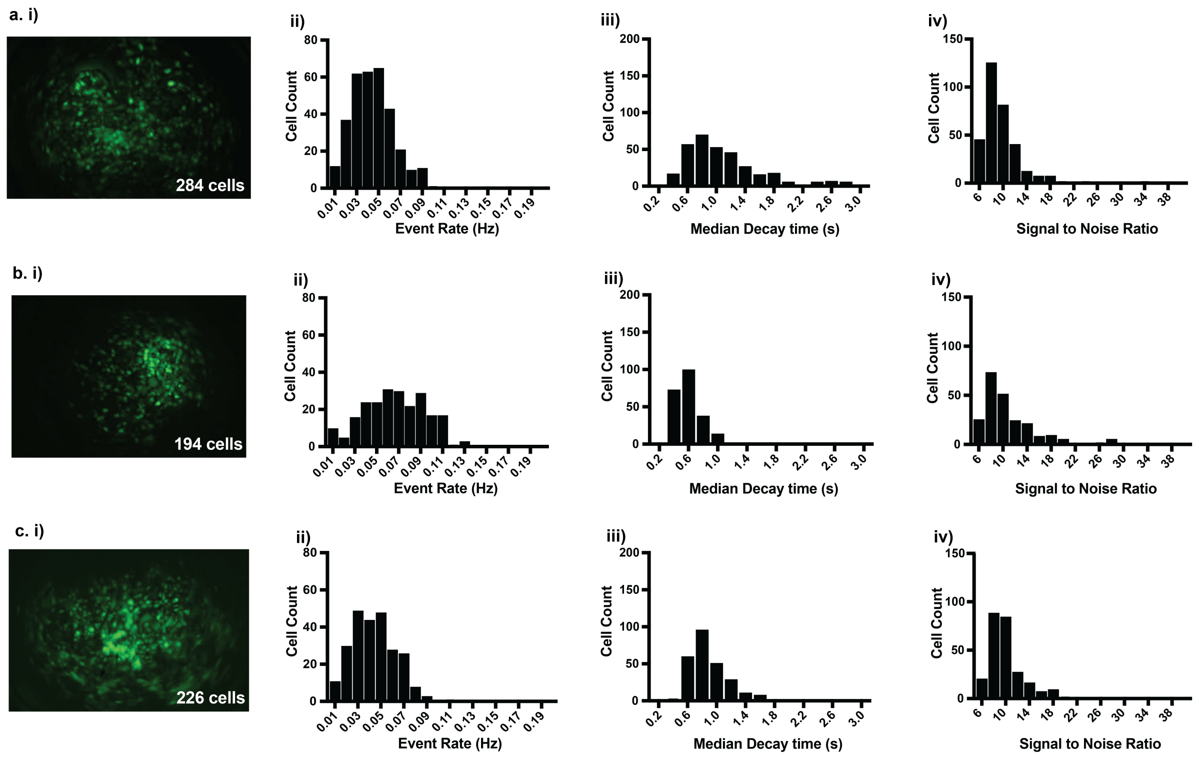


**Figure S6.** Replicates of fast implants of commercial silk films. **a)** First replicate of fast implant of silk film illustrating **i)** max projection image (GCaMP cells in green), histogram of neuronal calcium transient **ii)** median event rate, **iii)** median decay time, and **iv)** median signal to noise ratio. **b)** Second replicate of fast implant of silk film illustrating **i)** max projection image (GCaMP cells in green), histogram of neuronal calcium transient **ii)** median event rate, **iii)** median decay time, and **iv)** median signal to noise ratio. **c)** Third replicate of fast implant of silk film illustrating **i)** max projection image (GCaMP cells in green), histogram of neuronal calcium transient **ii)** median event rate, **iii)** median decay time, and **iv)** median signal to noise ratio.

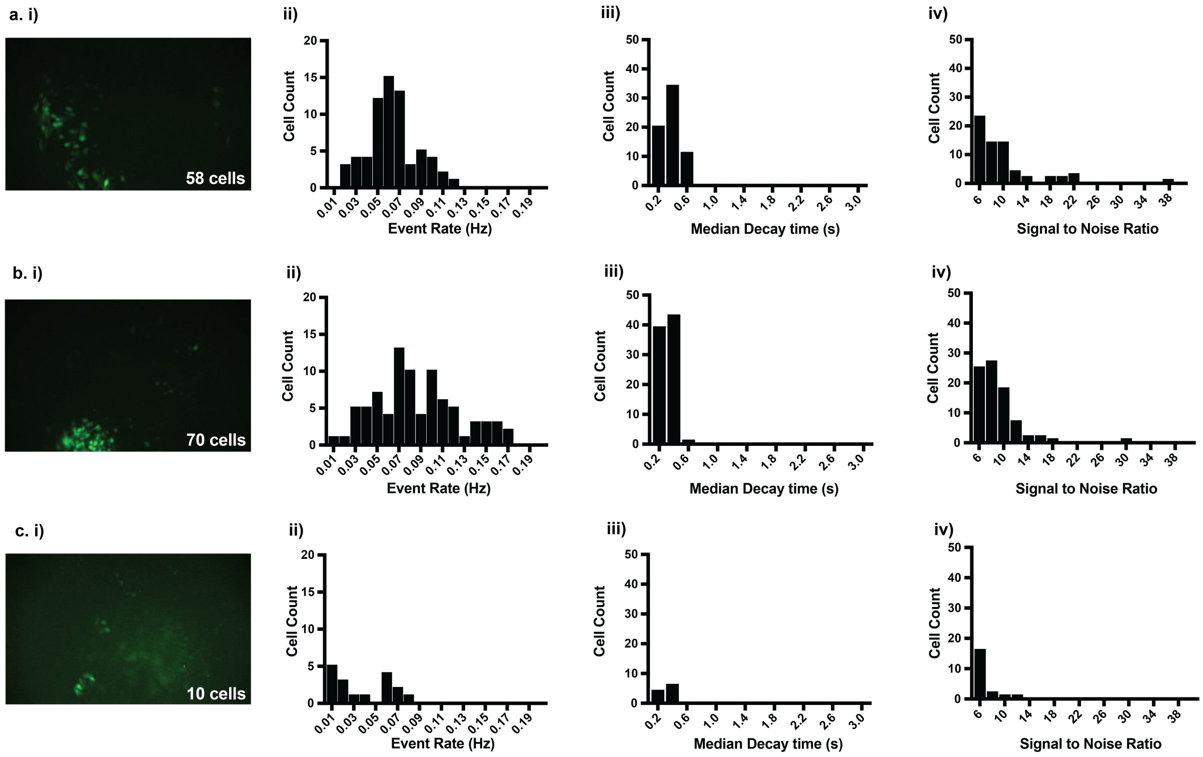


**Figure S7.** Replicates of slow implants of commercial silk films. **a)** First replicate of slow implant of silk film illustrating **i)** max projection image (GCaMP cells in green), histogram of neuronal calcium transient **ii)** medium event rate, **iii)** median decay time, and **iv)** median signal to noise ratio. **b)** Second replicate of slow implant of silk film illustrating **i)** max projection image (GCaMP cells in green), histogram of neuronal calcium transient **ii)** median event rate, **iii)** median decay time, and **iv)** median signal to noise ratio. **c)** Third replicate of slow implant of silk film illustrating **i)** max projection image (GCaMP cells in green), histogram of neuronal calcium transient **ii)** median event rate, **iii)** median decay time, and **iv)** median signal to noise ratio.

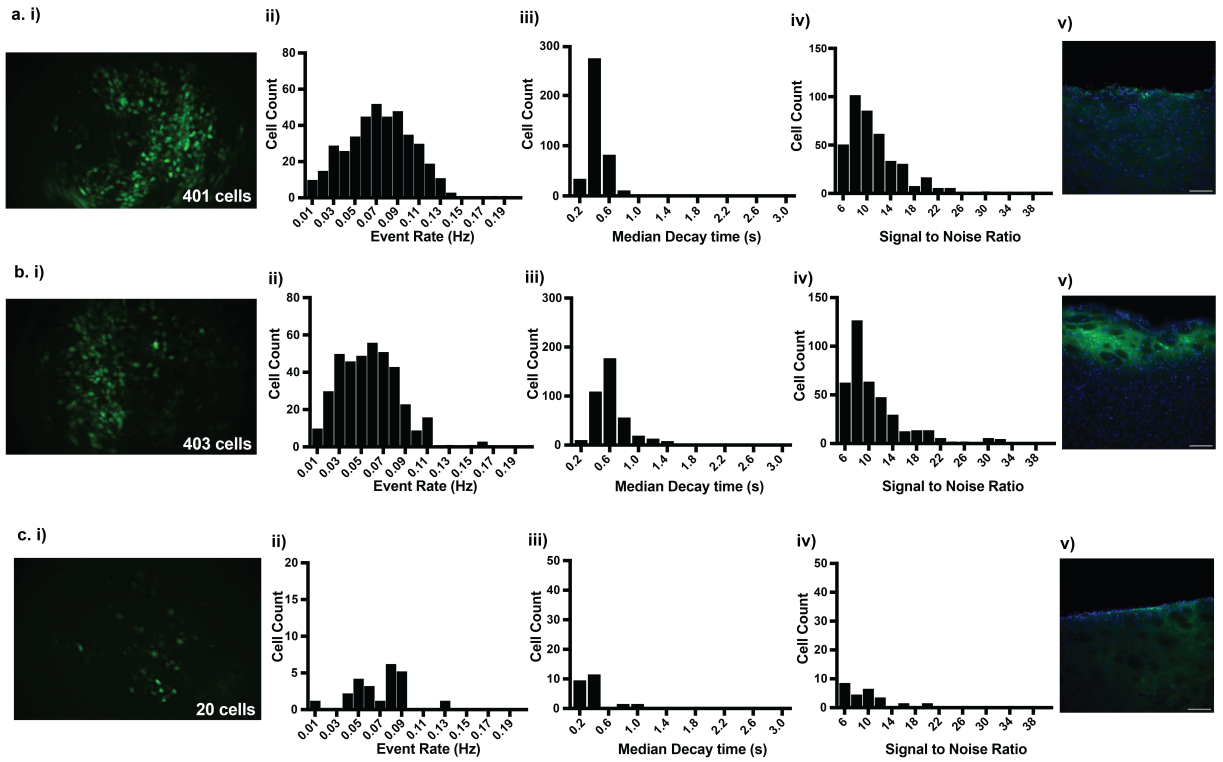


**Figure S8.** Replicates of fast and slow implants of CMC$_{30}$:Tre$_{70}$ films. **a)** First replicate of fast implant of CMC$_{30}$:Tre$_{70}$ film illustrating **i)** max projection image (GCaMP cells in green), histogram of neuronal calcium transient **ii)** median event rate, **iii)** median decay time, and **iv)** median signal to noise ratio, and **v)** post-mortem histology. **b)** Second replicate of fast implant of CMC$_{30}$:Tre$_{70}$ film illustrating **i)** max projection image (GCaMP cells in green), histogram of neuronal calcium transient **ii)** median event rate, **iii)** median decay time, and **iv)** median signal to noise ratio, and **v)** post-mortem histology. **c)** Replicate slow implant of CMC$_{30}$:Tre$_{70}$ film illustrating **i)** max projection image (GCaMP cells in green), histogram of neuronal calcium transient **ii)** median event rate, **iii)** median decay time, and **iv)** median signal to noise ratio, and **v)** post-mortem histology. Histology scale bars are 100 µm.

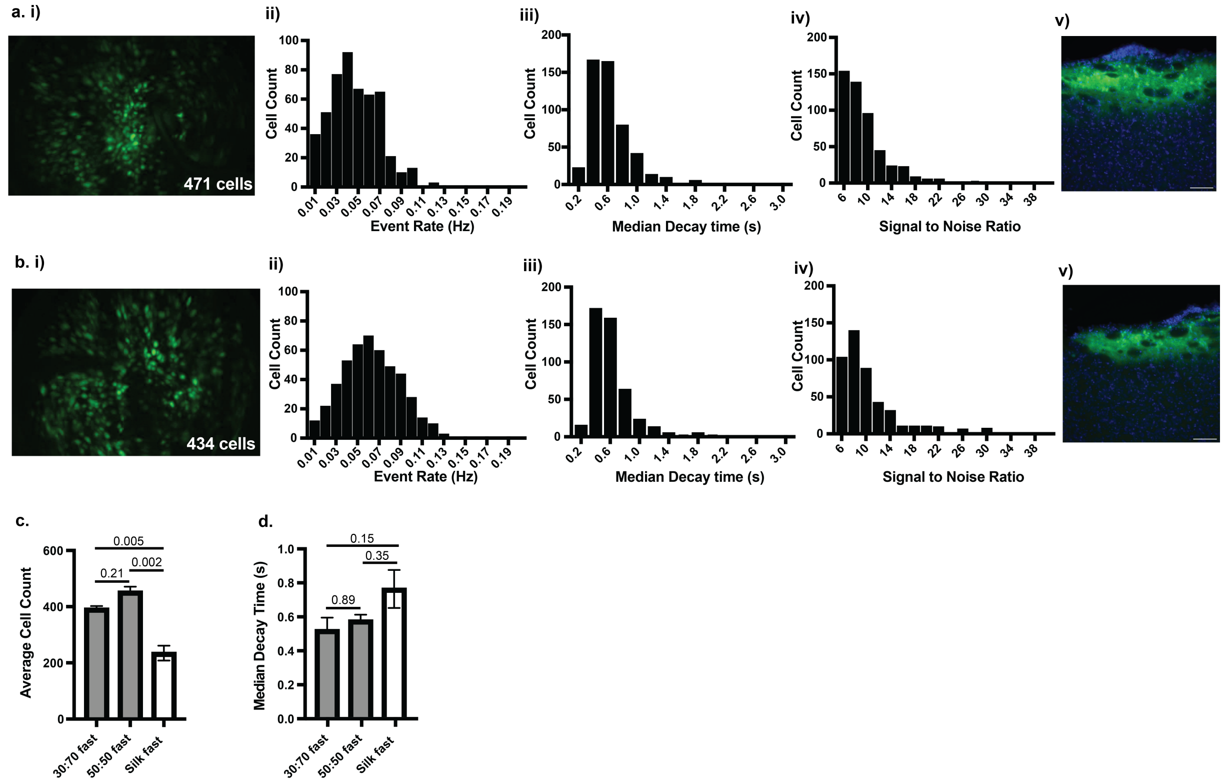


**Figure S9.** Comparative fast implants of CMC$_{50}$:Tre$_{50}$ films. **a)** First replicate of fast implant of CMC$_{50}$:Tre$_{50}$ film illustrating **i)** max projection image (GCaMP cells in green), histogram of neuronal calcium transient **ii)** median event rate, **iii)** median decay time, and **iv)** median signal to noise ratio, and **v)** post-mortem histology. **b)** Second replicate of fast implant of CMC$_{50}$:Tre$_{50}$ film illustrating **i)** max projection image (GCaMP cells in green), histogram of neuronal calcium transient **ii)** median event rate, **iii)** median decay time, and **iv)** median signal to noise ratio, and **v)** post-mortem histology. **c)** Comparison of average cell count for CMC$_{30}$:Tre$_{70}$ fast implants, CMC$_{50}$:Tre$_{50}$ fast implants, and silk fast implants. **d)** Comparison of median decay time for CMC$_{30}$:Tre$_{70}$ fast implants, CMC$_{50}$:Tre$_{50}$ fast implants, and silk fast implants. Histology scale bars are 100 µm. Values reported in plots c and d are means ± SEM of $n$ = 2-3 per group. Statistical significance values are $p$ values obtained from a Tukey HSD test.

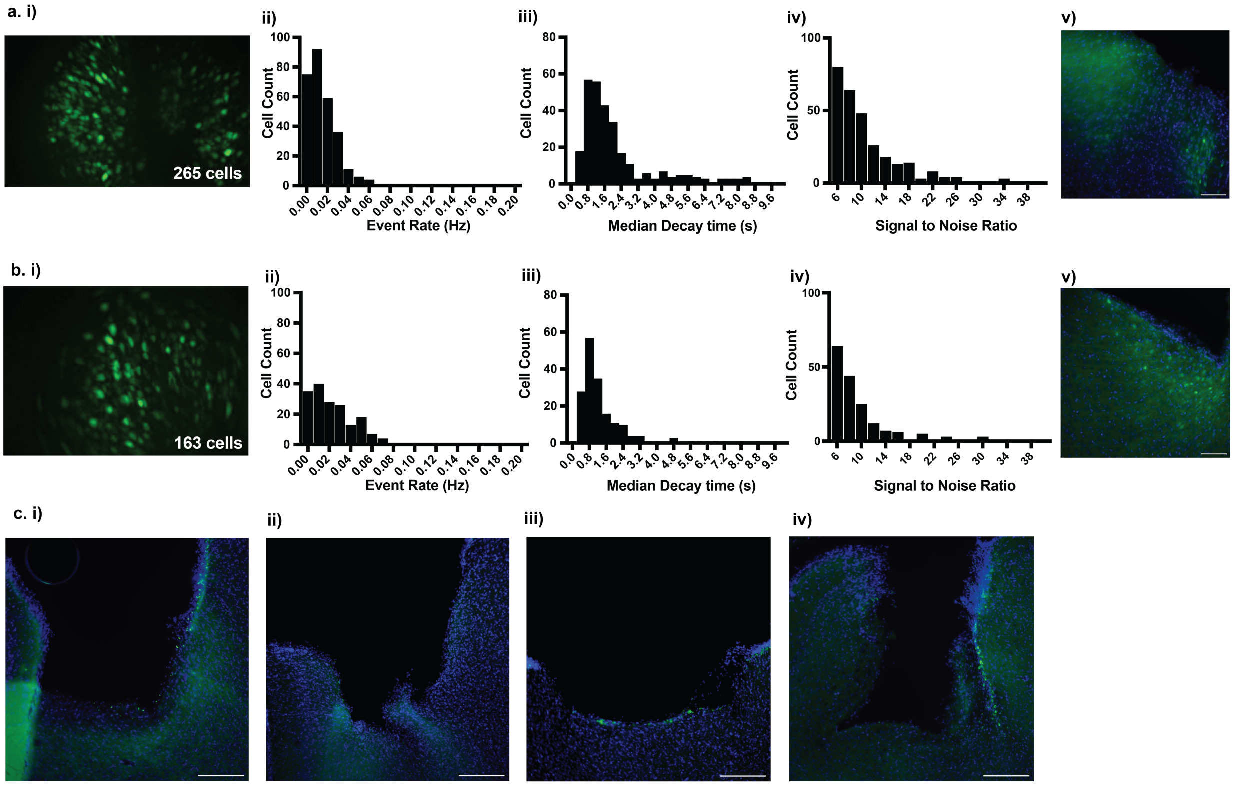


**Figure S10.** Replicates of fast and slow implants of CMC$_{30}$:Tre$_{70}$-G films. **a)** First replicate of fast implant of CMC$_{30}$:Tre$_{70}$-G film illustrating **i)** max projection image (GCaMP cells in green), histogram of neuronal calcium transient **ii)** median event rate, **iii)** median decay time, and **iv)** median signal to noise ratio, and **v)** post-mortem histology. **b)** Second replicate of fast implant of CMC$_{30}$:Tre$_{70}$-G film illustrating **i)** max projection image (GCaMP cells in green), histogram of neuronal calcium transient **ii)** median event rate, **iii)** median decay time, and **iv)** median signal to noise ratio, and **v)** post-mortem histology. **c)** Third replicate of fast implant of CMC$_{30}$:Tre$_{70}$-G film illustrating **i)** max projection image (GCaMP cells in green), histogram of neuronal calcium transient **ii)** median event rate, **iii)** median decay time, and **iv** median signal to noise ratio, and **v)** post-mortem histology. **d)** Fourth replicate of fast implant of CMC$_{30}$:Tre$_{70}$-G film illustrating **i)** max projection image (GCaMP cells in green), histogram of neuronal calcium transient **ii)** median event rate, **iii)** median decay time, and **iv)** median signal to noise ratio, and **v)** post-mortem histology. **e)** First replicate of slow implant of CMC$_{30}$:Tre$_{70}$-G film illustrating **i)** max projection image (GCaMP cells in green), histogram of neuronal calcium transient **ii)** median event rate, **iii)** median decay time, and **iv)** median signal to noise ratio, and **v)** post-mortem histology. **f)** Second replicate of slow implant of CMC$_{30}$:Tre$_{70}$-G film illustrating **i)** max projection image (GCaMP cells in green), histogram of neuronal calcium transient **ii)** median event rate, **iii)** median decay time, and **iv)** median signal to noise ratio, and **v)** post-mortem histology. Histology scale bars are 100 μm.

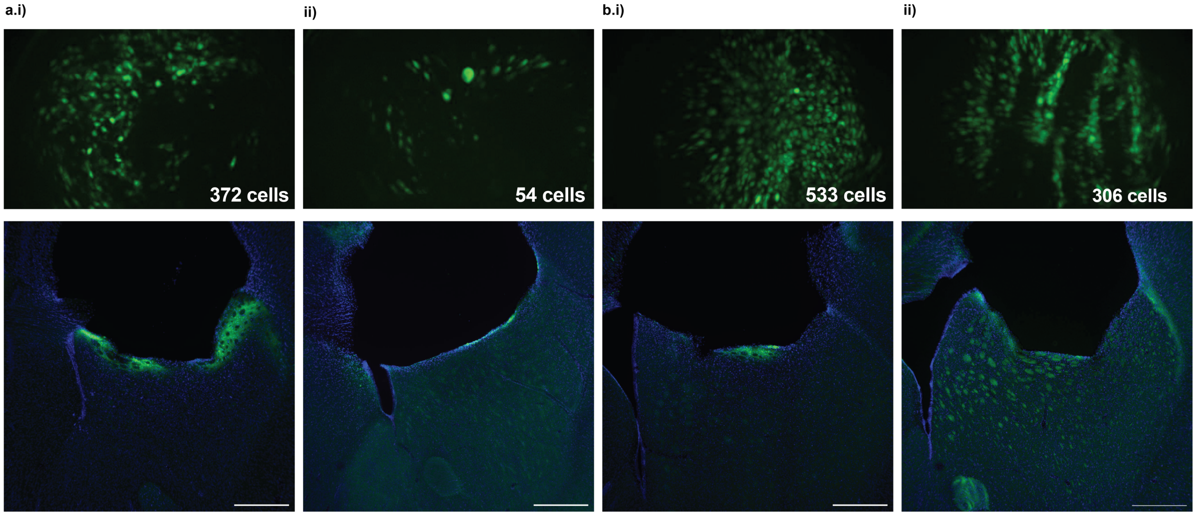


**Figure S11.** Insertion traces of fast and slow implants of $CMC_{30}$:$Tre_{70}$ and $CMC_{30}$:$Tre_{70}$-G films. **a)** Histology showing insertion trace and max projection image for **i)** fast implant and **ii)** slow implant of $CMC_{30}$:$Tre_{70}$ film. **b)** Histology showing insertion trace and max projection image for **i)** fast implant and **ii)** slow implant of $CMC_{30}$:$Tre_{70}$-G film. Histology scale bars are 250 µm.

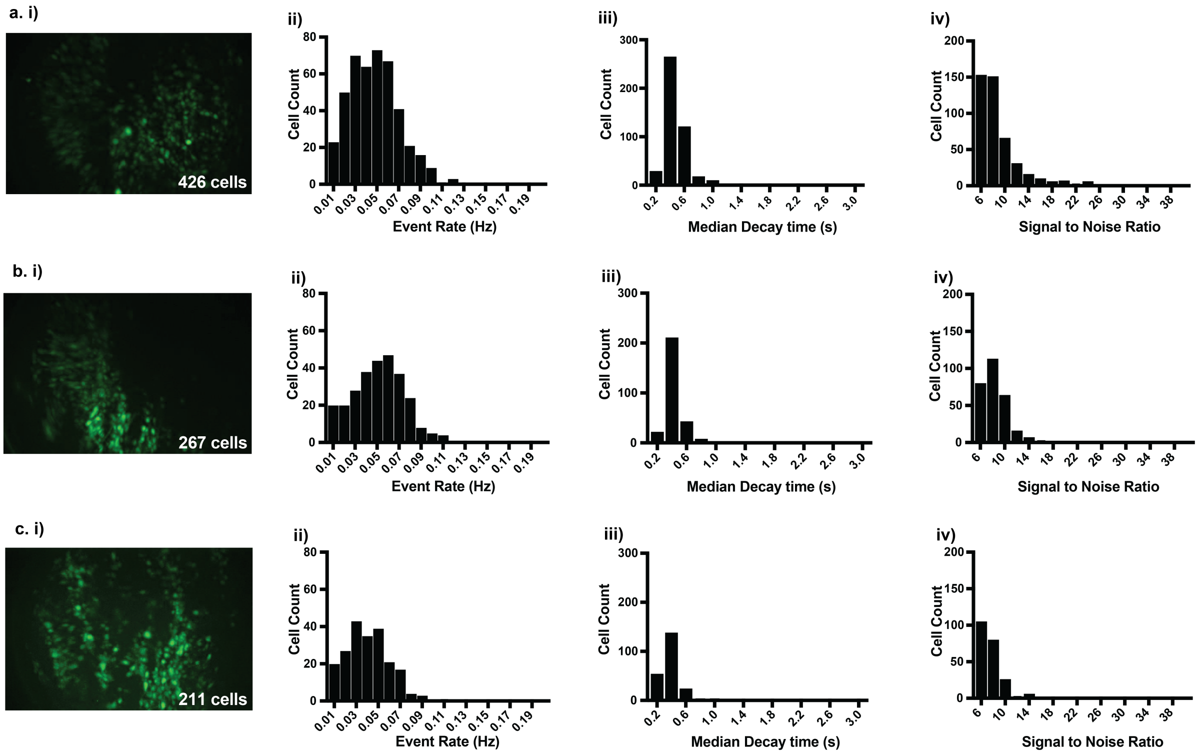


**Figure S12.** Replicates of fast implants of $CMC_{30}$:$Tre_{70}$-G films following 16 weeks of storage at 4C. **a)** First replicate of fast implant of stored $CMC_{30}$:$Tre_{70}$-G film illustrating **i)** max projection image (GCaMP cells in green), histogram of neuronal calcium transient **ii)** median event rate, **iii)** median decay time, and **iv)** median signal to noise ratio. **b)** Second replicate of fast implant of stored $CMC_{30}$:$Tre_{70}$-G film illustrating **i)** max projection image (GCaMP cells in green), histogram of neuronal calcium transient **ii)** median event rate, **iii)** median decay time, and **iv)** median signal to noise ratio. **c)** Third replicate of fast implant of stored $CMC_{30}$:$Tre_{70}$-G film illustrating **i)** max projection image (GCaMP cells in green), histogram of neuronal calcium transient **ii)** median event rate, **iii)** median decay time, and **iv)** median signal to noise ratio.

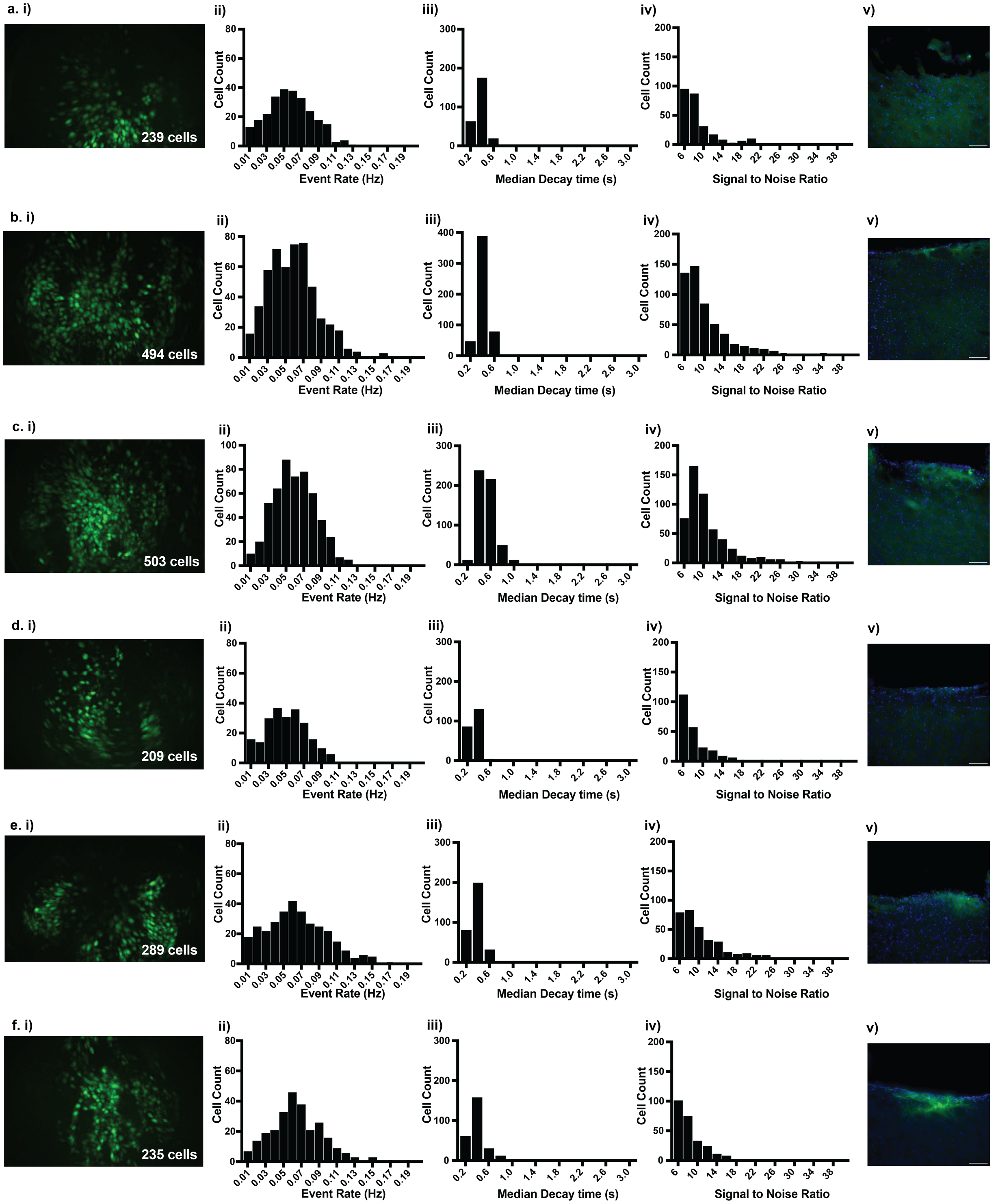


**Figure S13.** Replicates of $CMC_{30}$:$Tre_{70}$-G films implanted into the mPFC of mice and rats **a)** Replicate of $CMC_{30}$:$Tre_{70}$-G film implanted into the mPFC of a mouse illustrating **i)** max projection image (GCaMP cells in green), histogram of neuronal calcium transient **ii)** median event rate, **iii)** median decay time, and **iv)** median signal to noise ratio, and **v)** post-mortem histology. **b)** Replicate of $CMC_{30}$:$Tre_{70}$-G film implanted into the mPFC of a rat illustrating **i)** max projection image (GCaMP cells in green), histogram of neuronal calcium transient **ii)** median event rate, **iii)**

median decay time, and **iv)** median signal to noise ratio, and **v)** post-mortem histology. Histology scale bars in v. are 100 µm. **c)** Post-mortem histology of replicates of commercial silk film implants **(i, ii, iii, iv)** in the mouse mPFC. Implants illustrate little to no GCaMP expression below the imaging lens. Histology scale bars in c are 250 um. Max projection images and histograms of neuronal calcium transient event rates, decay times, and signal to noise ratio are not included because commercial silk film implants resulted in no identifiable GCaMP expressing cells.

**Table S1.** Summary of probe coating and implant procedure as well as corresponding total cells, event rate, decay time, and signal to noise ratio.

| Coatings | Implant Procedure | Cells in Field of View | | Median Event Rate (Hz) | | Median Decay Time (sec) | | Median Signal to Noise Ratio | |
|---|---|---|---|---|---|---|---|---|---|
| | | | Avg | | Avg | | Avg | | Avg |
| CMC$_{30}$:Tre$_{70}$ | mouse dorsal striatum, fast | 372 | 392 | 0.044 | 0.057 | 0.5 | 0.52 | 8.24 | 8.95 |
| CMC$_{30}$:Tre$_{70}$ | mouse dorsal striatum, fast | 403 | | 0.054 | | 0.6 | | 8.83 | |
| CMC$_{30}$:Tre$_{70}$ | mouse dorsal striatum, fast | 401 | | 0.072 | | 0.45 | | 9.77 | |
| CMC$_{30}$:Tre$_{70}$ | mouse dorsal striatum, slow | 54 | 37 | 0.049 | 0.062 | 0.4 | 0.38 | 7.19 | 8.05 |
| CMC$_{30}$:Tre$_{70}$ | mouse dorsal striatum, slow | 20 | | 0.075 | | 0.35 | | 8.90 | |
| CMC$_{50}$:Tre$_{50}$ | mouse dorsal striatum, fast | 434 | 453 | 0.061 | 0.053 | 0.55 | 0.58 | 8.70 | 8.51 |
| CMC$_{50}$:Tre$_{50}$ | mouse dorsal | 471 | | 0.045 | | 0.60 | | 8.32 | |

| | | | | | | | | | |
|---|---|---|---|---|---|---|---|---|---|
| | striatum, fast | | | | | | | | |
| $CMC_{30}$:$Tre_{70}$-G | mouse dorsal striatum, fast | 533 | 396 | 0.057 | 0.055 | 0.50 | 0.45 | 9.29 | 8.33 |
| $CMC_{30}$:$Tre_{70}$-G | mouse dorsal striatum, fast | 239 | | 0.055 | | 0.38 | | 7.65 | |
| $CMC_{30}$:$Tre_{70}$-G | mouse dorsal striatum, fast | 494 | | 0.056 | | 0.45 | | 8.44 | |
| $CMC_{30}$:$Tre_{70}$-G | mouse dorsal striatum, fast | 503 | | 0.058 | | 0.55 | | 9.23 | |
| $CMC_{30}$:$Tre_{70}$-G | mouse dorsal striatum, fast | 209 | | 0.049 | | 0.35 | | 7.02 | |
| $CMC_{30}$:$Tre_{70}$-G | mouse dorsal striatum, slow | 306 | 277 | 0.099 | 0.075 | 0.40 | 0.39 | 8.42 | 8.63 |
| $CMC_{30}$:$Tre_{70}$-G | mouse dorsal striatum, slow | 289 | | 0.063 | | 0.38 | | 8.87 | |
| $CMC_{30}$:$Tre_{70}$-G | mouse dorsal striatum, slow | 235 | | 0.063 | | 0.38 | | 7.59 | |
| silk fibroin | mouse dorsal striatum, fast | 284 | 235 | 0.044 | 0.051 | 0.94 | 0.76 | 8.71 | 9.13 |
| silk fibroin | mouse dorsal | 194 | | 0.067 | | 0.55 | | 9.46 | |

| | | | | | | | | | |
|---|---|---|---|---|---|---|---|---|---|
| | striatum, fast | | | | | | | | |
| silk fibroin | mouse dorsal striatum, fast | 226 | | 0.042 | | 0.80 | | 9.21 | |
| silk fibroin | mouse dorsal striatum, slow | 58 | 46 | 0.061 | 0.067 | 0.38 | 0.36 | 8.75 | 7.79 |
| silk fibroin | mouse dorsal striatum, slow | 70 | | 0.080 | | 0.35 | | 8.21 | |
| silk fibroin | mouse dorsal striatum, slow | 10 | | 0.059 | | 0.34 | | 6.42 | |
| CMC$_{30}$:Tre$_{70}$-G | mouse mPFC | 283 | 274 | 0.017 | 0.015 | 1.25 | 1.38 | 9.80 | 9.29 |
| CMC$_{30}$:Tre$_{70}$-G | mouse mPFC | 265 | | 0.012 | | 1.50 | | 8.78 | |
| CMC$_{30}$:Tre$_{70}$-G | rat mPFC | 124 | 144 | 0.007 | 0.012 | 0.95 | 0.98 | 6.93 | 7.38 |
| CMC$_{30}$:Tre$_{70}$-G | rat mPFC | 163 | | 0.017 | | 1.0 | | 7.82 | |

**Video S1.** CMC$_{30}$:Tre$_{70}$-G fast implant into the mouse dorsal medial striatum

https://drive.google.com/file/d/1jzIEZyg7t3SyVRqKPRHMQAIfjtVUiyUs/view?usp=share_link

**Video S2.** CMC$_{30}$:Tre$_{70}$-G slow implant into the mouse dorsal medial striatum

https://drive.google.com/file/d/1igbvXfAOKlnIBVVMTVKG10kPmMpSaOIR/view?usp=share_link

**Video S3.** CMC$_{30}$:Tre$_{70}$-G implant into the mouse prefrontal cortex

https://drive.google.com/file/d/1BvZVe9FgxyCitu12oyS1wIDXQTgvaioQ/view?usp=share_link

**Video S4.** CMC$_{30}$:Tre$_{70}$-G implant into the rat prefrontal cortex https://drive.google.com/file/d/1fW84HnW2boJouTyYhqqmWZaeBXHNNZYs/view?usp=share_link
